# Nucleus-level thalamic organization anchors multimodal signatures of thalamocortical maturation

**DOI:** 10.64898/2026.07.03.736332

**Authors:** Alexandra John, Amin Saberi, Aikaterina Manoli, Jessica Royer, Doruk Yigit Eriguec, Valerie J. Sydnor, Bin Wan, Simon B. Eickhoff, Boris C. Bernhardt, Alfred Anwander, Sofie L. Valk

## Abstract

The human thalamus is composed of multiple nuclei that differ in structure and function. From early development onwards, these nuclei form reciprocal, nucleus-specific connections with the cerebral cortex, contributing to sensory and cognitive processing. In childhood and adolescence, a key period of neurocognitive development, these connections undergo widespread refinement, yet how developmental trajectories of thalamocortical connections vary across nuclei remains unknown. Here, we leveraged the Human Connectome Project in Development dataset (HCP-D, *N* = 604, age range 8-21) and segmented 10 thalamic nuclei using a segmentation approach optimized for intrathalamic contrast. Applying probabilistic tractography, we reconstructed nucleus-specific thalamocortical connections and charted their maturational profiles based on changes in fractional anisotropy (FA) using generalized additive models. We found FA to increase in thalamocortical connections, with nucleus-specific variation in temporal profiles and magnitude of age effects. Connections of core-cell-rich, sensory-projecting nuclei, such as the lateral geniculate nucleus, showed earlier maturational plateaus, whereas matrix-cell-rich, association-projecting nuclei, such as ventral anterior nucleus, showed more sustained maturation. This links maturational heterochronicity to thalamic organization of cell distribution and connectivity embedding. In parallel, functional thalamocortical connectivity decreased with age, with FA and functional connectivity age effects coupled in nucleus-connections showing prolonged maturation. Finally, concordant age effects in connectivity and nucleus volumes suggest that intra-nucleus remodeling may support refinement of structural connections while reducing thalamocortical functional synchrony. Together, our work reveals that thalamocortical maturation is anchored in the developmental and organizational heterogeneity of thalamic nuclei, offering a framework for understanding how diverse thalamic nuclei contribute to neurocognitive development.

## Introduction

The thalamus is a centrally located, subcortical hub within large-scale brain organization, forming extensive bidirectional connections with the entire cerebral cortex. These widespread thalamocortical connections enable the thalamus to strongly impact cortical dynamics and function (Shine et al., 2023; Hwang et al., 2017; Malekmohammadi et al., 2015; Nakajima and Halassa, 2017; Bagshaw, 2026). Akin to the structural and functional diversity of regions within the cerebral cortex, the thalamus comprises a highly heterogeneous architecture of multiple nuclei with specialized roles in sensory processing and cognition (Shine et al., 2023; Wolff et al., 2021). Thalamic nuclei originate from different progenitors, vary markedly in their cytoarchitectural composition, and exhibit unique nucleus-specific cortical connectivity profiles (Govek et al., 2022; Kim et al., 2023; Lo Giudice et al., 2024; Morel et al., 1997; Clascá et al., 2012; Lambert et al., 2017). These thalamocortical connections are established pre- and early postnatally (Wilson et al., 2023; Grant et al., 2012), but like the rest of the brain, undergo continuous changes throughout development (Oldham et al., 2025; Sydnor et al., 2025; Zheng et al., 2023). An important developmental period in the brain is the phase of childhood and adolescence, which is marked by heightened cortical plasticity supporting the refinement and emergence of cognitive functions (Baum et al., 2020; Larsen and Luna, 2018; Arain et al., 2013). Prior work suggests that the development of thalamocortical connections in childhood and adolescence mirrors cortical development and may help orchestrate windows of cortical plasticity (Sydnor et al., 2025). However, despite the pronounced heterogeneity of thalamic nuclei, their nucleus-specific developmental trajectories remain unresolved, limiting our ability to relate thalamocortical maturation to thalamic organization and development. Here, we ask how individual nuclei develop in their connectivity with the cortex, and how this relates to intrathalamic organization, functional connectivity and nucleus volume maturation, providing a thalamocentric account of brain maturation in childhood and adolescence.

The bidirectional information flow between thalamic nuclei and their cortical targets is mediated by reciprocal thalamocortical connections, allowing thalamic and cortical regions to dynamically modulate one another (Hwang et al., 2022; Shine et al., 2023; Briggs and Usrey, 2008; Crandall et al., 2015; Sherman, 2005, 2016; Malekmohammadi et al., 2015). Therefore, thalamic nuclei are no longer considered as passive ascending relay stations. Instead, they actively contribute to many functions, ranging from sensory processing in nuclei such as the lateral geniculate nucleus (LGN) and medial geniculate nucleus (MGN) to more abstract cognitive processes, such as attention and cognitive control in the medial dorsal nucleus (MD) and Pulvinar (Saalmann and Kastner, 2011; Zhou et al., 2016; Saalmann et al., 2012; Rikhye et al., 2018; Ferguson and Delevich, 2020). Beyond this, thalamic nuclei regulate cortical activity and support the integration and segregation of cortical regions by linking them through trans-thalamic connections (Shine et al., 2023; Malekmohammadi et al., 2015; Sherman and Guillery, 2011). The functional variety of thalamic nuclei is reflected in partially overlapping organizational characteristics, spanning differences in cortical projection patterns, cellular architecture, and input sources. Structural connections between thalamic nuclei and the cortex vary substantially in their cortical target areas, cortical layers, and the degree of how diffusely or selectively they project. The differences in connectivity patterns are closely linked to the distribution of neuronal core- and matrix cells within the thalamus (Clascá et al., 2012; Jones, 1998; Müller et al., 2020). Sensory thalamic nuclei tend to have a higher proportion of core cells, which project preferentially in a targeted manner to cortical layers IV and V, whereas matrix cells are characterized by distributed cortical projections into layer I-III (Jones, 1998; Clascá et al., 2012; Müller et al., 2020). Another characterization of thalamic nuclei is based on the origin of the main ‘driver’ input. A distinction is made between first order nuclei that receive input primarily from the subcortex and higher order nuclei that receive main driver input from cortical layer V (Sherman, 2005, 2016). Together, the differences in microstructure, function, and connectivity underscore the heterogeneity of thalamic nuclei and highlight the importance of studying them as distinct structures.

The strong interaction between the thalamic nuclei and the cortex is established in early development. While initial cortical arealization (including neurogenesis and neuronal migration) is largely governed by intrinsic genetic factors, once thalamocortical connections are formed prenatally, the thalamus and cortex shape each other’s maturation (Molnár and Kwan, 2024; Antón-Bolaños et al., 2018; Rakic, 1988). Insights from animal models have shown that thalamic cues are crucial for cortical maturation, such as the definition of cortical area size and identity (Antón-Bolaños et al., 2018), sensory topography (*e.g.,* formation of barrel organization in mouse sensory cortex) (Molnár and Kwan, 2024), the development of inhibitory interneurons (De Marco García et al., 2015), and cortical laminar development (Sato et al., 2022; Monko et al., 2022). Vice versa, corticofugal activity impacts the development of thalamocortical connections and thalamic nuclei, such as axon targeting and nucleus size (Antón-Bolaños et al., 2018; Molnár and Kwan, 2024; Li et al., 2023; Thompson et al., 2016). This interplay in early development underlines the co-dependent formation of thalamic nuclei and cortex as an interlinked circuit, coordinated by reciprocal thalamocortical connections.

The maturation of the thalamocortical system continues throughout childhood and adolescence. As this developmental period is crucial for the refinement of cognitive abilities, such as executive function and social cognition, and marked by increased vulnerability to neuropsychiatric disorders (Larsen and Luna, 2018; Sydnor et al., 2021; Larsen et al., 2023; Park et al., 2022; Baum et al., 2017), understanding the underlying spatiotemporal patterns of brain maturation across heterogeneous regions is essential. Yet, studies addressing this question have primarily focused on the cerebral cortex, where a growing body of evidence revealed a hierarchical spatiotemporal development along the cortical sensorimotor-association (SA) axis. Along this axis, sensorimotor areas mature earlier, whereas association areas show a prolonged time window of maturation (Sydnor et al., 2021, 2023; Luo et al., 2024; Taylor et al., 2026; Dong et al., 2021), with cortical myelination providing one example following this pattern (Baum et al., 2022; Paquola et al., 2019). An important advance in understanding how cortical development relates to thalamocortical connections was provided by Sydnor *et al*., 2025 (Sydnor et al., 2025). This study showed that the heterochronous maturation of thalamocortical white matter connection parallels the SA axis of cortical development, with earlier maturation in sensory connections and prolonged maturation in connections to association regions. The maturation of structural connections was measured by changes in fractional anisotropy (FA), a metric that quantifies the directionality of water diffusion within a voxel. While FA is sensitive to different biophysical aspects, in development, increasing FA is attributed to axon myelination, which suggests more efficient neural transmission (Paus, 2010). Complementing neuroimaging studies, animal work supports that thalamic circuits undergo experience-dependent sculpting in adolescence (Nakayama and Miyata, 2025). Together, the refinement of thalamocortical connections is central to neurodevelopment in childhood and adolescence. However, despite the heterogeneity of thalamic nuclei in structure, function, and connectivity patterns, their individual contributions to this process and their interrelations are unknown.

Of note, emerging evidence suggests that the heterogeneity of thalamic nuclei extends to their developmental trajectories and vulnerability in neurodevelopmental disorders, strengthening the rationale of moving beyond a whole-thalamus towards a nucleus-level perspective to understand thalamocortical maturation. For example, studies show an overall volume decrease of the whole-thalamus during childhood and adolescence, however maturational trajectories of individual nuclei differ, with some showing a volume decrease, while other areas expand (Carrick et al., 2025; Sussman et al., 2016; Huang et al., 2024; Herting et al., 2018; Raznahan et al., 2014). Similarly, whole-thalamus thalamocortical functional connectivity was shown to decrease overall, but again this is not uniform across nuclei (Steiner et al., 2020). Moreover, the heterogeneity among thalamic nuclei is reflected in neurodevelopmental disorders, where distinct nuclei and their cortical connections are differentially involved, including schizophrenia (Thalhammer et al., 2024b; Young et al., 2025; Huang et al., 2024, 2020; Pergola et al., 2015; Tanrıseven et al., 2026), autism spectrum conditions (Nair et al., 2013; Cerliani et al., 2015), as well as focal and generalized epilepsy (Lindquist et al., 2023; He et al., 2015). These findings motivate the investigation of nucleus-level thalamocortical structural connectivity maturation and its relation to thalamic organization, functional connectivity, and nucleus volume development. However, resolving structural connections at the level of individual nuclei has been challenging due to the small size, the close spatial proximity of the nuclei, and limitations of current neuroimaging techniques.

The present study navigates these challenges to examine how nucleus-specific thalamocortical connections mature in childhood and adolescence. By leveraging a large cross-sectional dataset, we reconstructed individualized nucleus-specific thalamocortical structural connections using probabilistic tractography followed by multistage filtering in order to remove spurious or unreliable connections. Next, we assessed the developmental trajectories of microstructural changes within these nucleus-specific connections in childhood and adolescence. This nucleus-resolved approach enabled us to test the hypothesis that developmental patterns of nucleus-specific thalamocortical structural connections are systematically related to intrinsic thalamic features. Moreover, we leveraged a multimodal approach and investigated how developmental changes in nucleus-specific thalamocortical structural connectivity are related to developmental changes in thalamocortical functional connectivity and nucleus volume development. In doing so, we move beyond a whole-thalamus perspective by acknowledging the heterogeneity of thalamic nuclei and their cortical connections, and characterize their contribution to brain development in childhood and adolescence.

## Results

To chart the developmental trajectories of nucleus-specific thalamocortical white matter connections during childhood and adolescence and investigate their link to multimodal thalamic features, we leveraged multimodal data from a cross-sectional sample (HCP-D, *N* = 604, age range 8-21, **Supplementary Fig. 1**). The thalamus was segmented into 10 nuclei at the individual level using the state-of-the-art ‘histogram-based polynomial synthesis - thalamus optimized multi-atlas’ approach (HIPS-THOMAS) (Vidal et al., 2024), which enhances intrathalamic contrast and is histologically-informed by the Morel atlas (Morel et al., 1997). Thalamocortical connections were reconstructed using probabilistic tractography, using thalamic nuclei as seeds and cortical Glasser parcels (Glasser et al., 2016) as target regions. Within the identified connections, FA, a measure of white matter microstructure that captures the degree of diffusion anisotropy (Lebel and Beaulieu, 2011; Lebel and Deoni, 2018), was quantified. Following, age-related effects of FA were modeled using generalized additive models (GAMs), while controlling for sex and head motion. We first assessed developmental patterns of whole-thalamus connectivity as a validation step, to test whether our tractography and modeling approaches reproduce previously reported findings of hierarchical thalamocortical connection development (Sydnor et al., 2025) (**Fig. 1**). Building on this, we then charted age-related developmental trajectories of thalamocortical connections at the level of individual thalamic nuclei (**Fig. 2**), and investigated their relation to nucleus-specific thalamic cell type composition, the categorization into first- and higher order nuclei, and connectivity embedding along the SA axis (**Fig. 3**), age-related changes of functional connectivity (**Fig. 4**), and corresponding thalamic nuclei development, indexed by volume changes (**Fig. 5**).

**Fig. 1:**
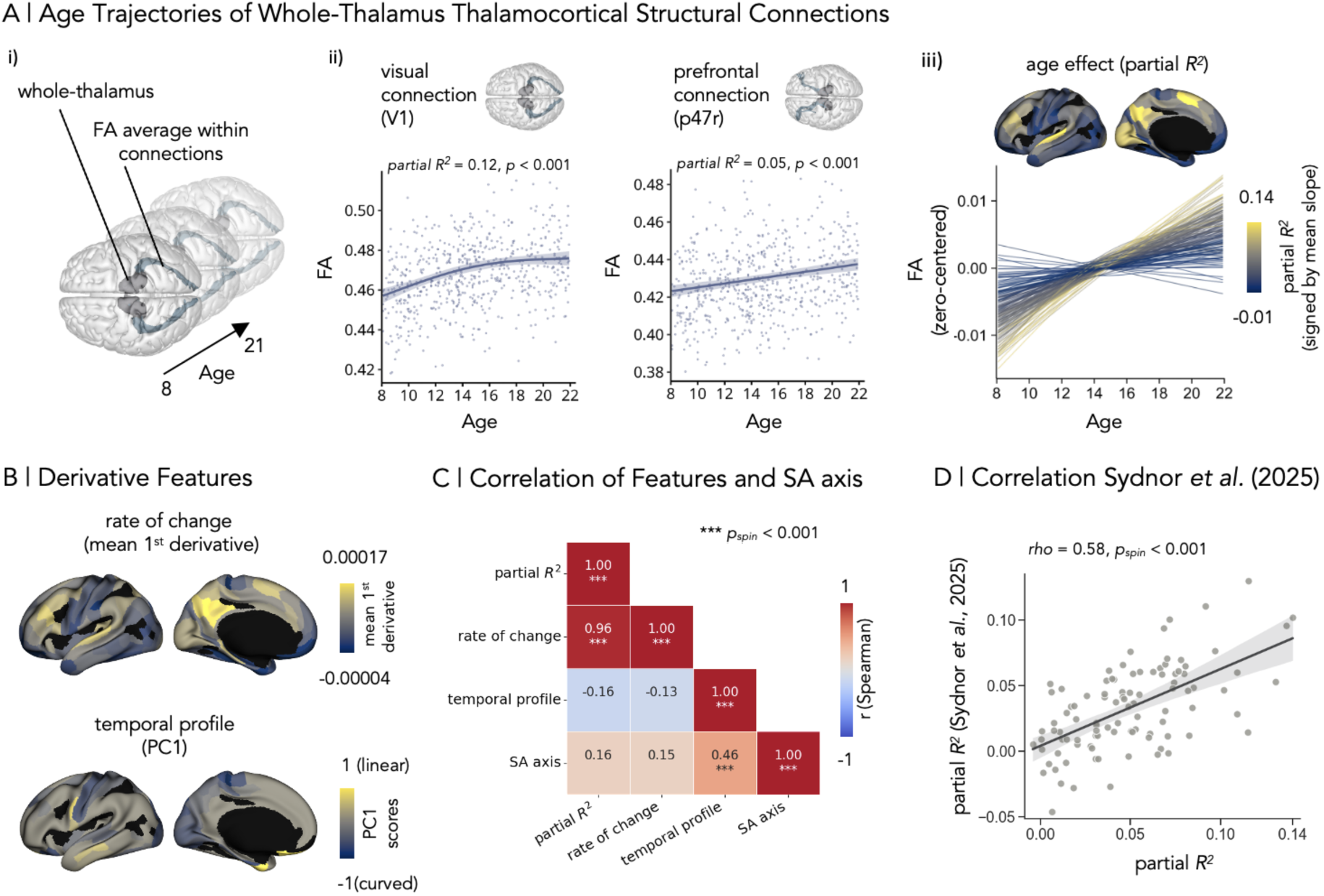
Developmental trajectories of whole-thalamus thalamocortical connectivity. **A** Developmental trajectories of whole-thalamus thalamocortical connections. i) Schematic representation showing the connection from the whole-thalamus to visual cortex (V1). Fractional anisotropy (FA) average within the tracts was extracted for all subjects (*N* = 604; age range 8–21). FA values of corresponding connections in left and right hemispheres were averaged for all follow-up analysis. ii) Age trajectories of FA from whole-thalamus-to-visual cortex (left subpanel) and whole-thalamus-to-prefrontal cortex (right subpanel), overlaid on individual FA data points. Trajectories show predicted FA values and the 95% confidence interval based on generalized additive models (GAMs). iii) Developmental GAM trajectories of all whole-thalamus thalamocortical connections. The trajectories represent zero-centered GAM smooth estimates and are colored by partial *R^2^*, which describes the magnitude of age effect and is signed by mean slope. The distribution of age effects are shown projected on the cortex by coloring the connections target parcel by partial *R^2^*. **B** Rate of change was calculated by the first derivative along the age trajectories and principal component analysis (PCA) was applied to the normalized first derivatives sampled along the trajectory across trajectories, indicating temporal profiles of FA change. Mean rate of change across age and the principal component 1 (PC1), which is referred to as temporal profile component, scores were projected on the cortex. **C** Correlation matrix showing Spearman correlations between partial *R^2^*, rate of change (mean first derivative), temporal profile component (PC1), and the sensorimotor-to-association (SA) axis (Sydnor et al., 2021). Spin permutation was applied and *p*_spin_ < 0.001 are indicated by ‘***’. The spatiotemporal pattern of the SA axis is shown in Fig 3. **D** Connection-wise partial *R^2^* from this study was correlated (Spearman) with connection-wise partial *R^2^* from Sydnor *et al*. (2025) (linear regression line with a shaded 95% confidence interval).

**Fig. 2:**
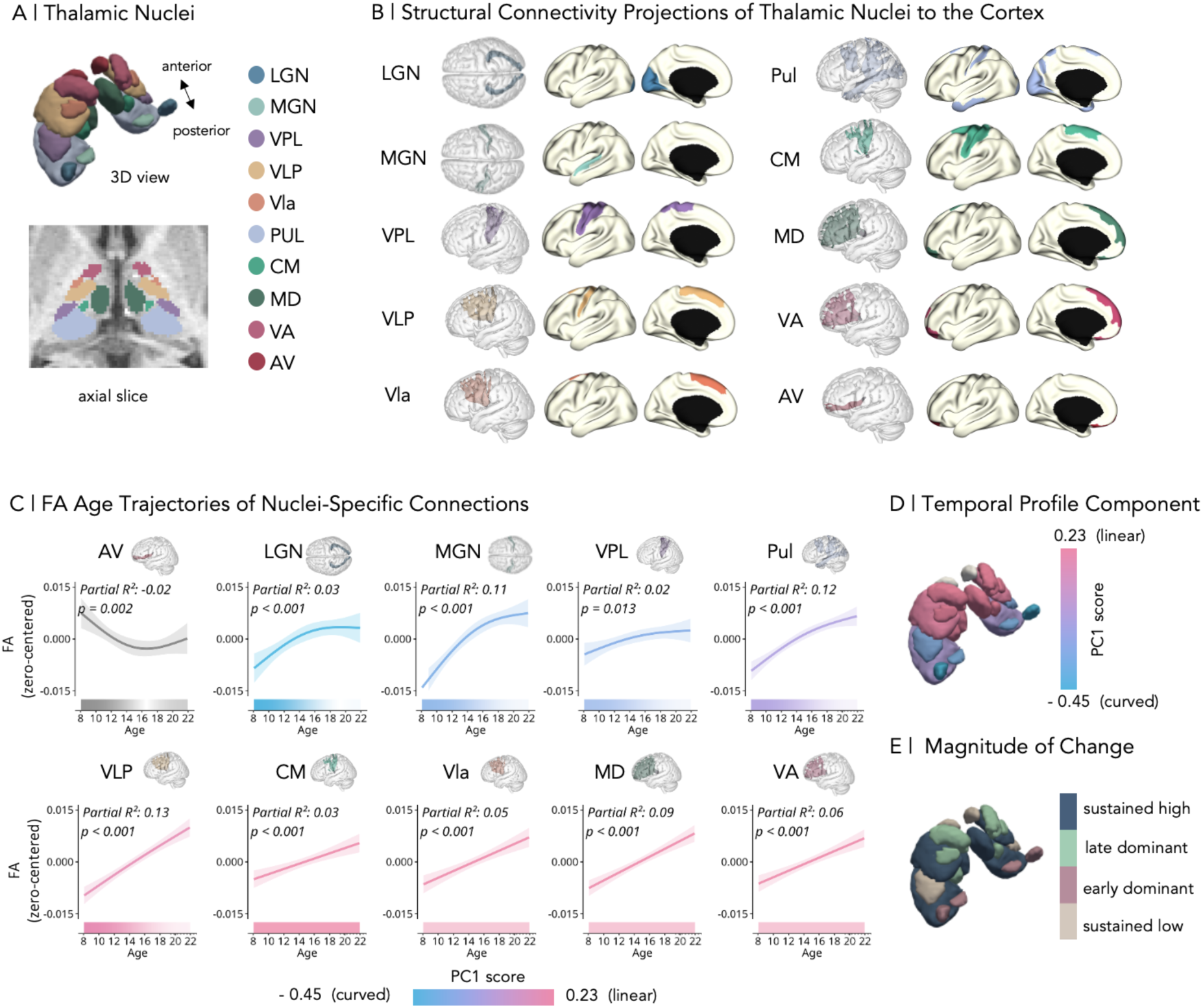
Developmental trajectories of nucleus-specific thalamocortical connections. **A** Thalamic nuclei segmentation shown as 3D rendering and projected on an axial plane of the thalamus (one example subject). **B** Nucleus-specific thalamocortical structural connections. For each nucleus, the thalamocortical connection (one example subject in native space) and the target parcels (generated at group-level) are mapped on the cortical surface. Maps are shown for the left hemisphere only. Right hemisphere results are shown in **Supplementary Fig. 3**. **C** Fractional anisotropy (FA)-age trajectories of cortical connections for each nucleus (FA of left and right hemisphere was averaged). The trajectories represent zero-centered generalized additive model (GAM)-smooth estimates with a shaded 95% confidence interval and are colored and ordered by principal component 1 score (PC1), referred to as temporal profile component (see **D**). AV was excluded from subsequent analysis due to insufficient connectivity reconstruction, and is therefore shown in grey. Partial *R²* (signed by mean slope), and *p*-value (FDR-corrected) are indicated. Shaded bars depict the rate of change measured by the first derivative. **D** PCA was applied to normalized first derivatives sampled along nucleus-specific age trajectories (AV excluded) and temporal profile scores were projected on the thalamus. **E** Magnitude of change projected on the thalamus. The mean first derivative of each nucleus was compared to the average change across all nuclei across the whole age range (sustained high, sustained low) and within the first half (early dominant) and second half (late dominant) of the age range. Abbreviations: Lateral Geniculate Nucleus (LGN), Medial Geniculate Nucleus (MGN), Ventral Posterior Lateral (VPL), Ventral Lateral Nucleus (posterior) (VLP), Ventral Lateral Nucleus (anterior) (VLa), Pulvinar (Pul), Centromedian Nucleus (CM), Mediodorsal Nucleus (MD), Ventral Anterior Nucleus (VA), Antero-ventral Nucleus (AV).

**Fig. 3:**
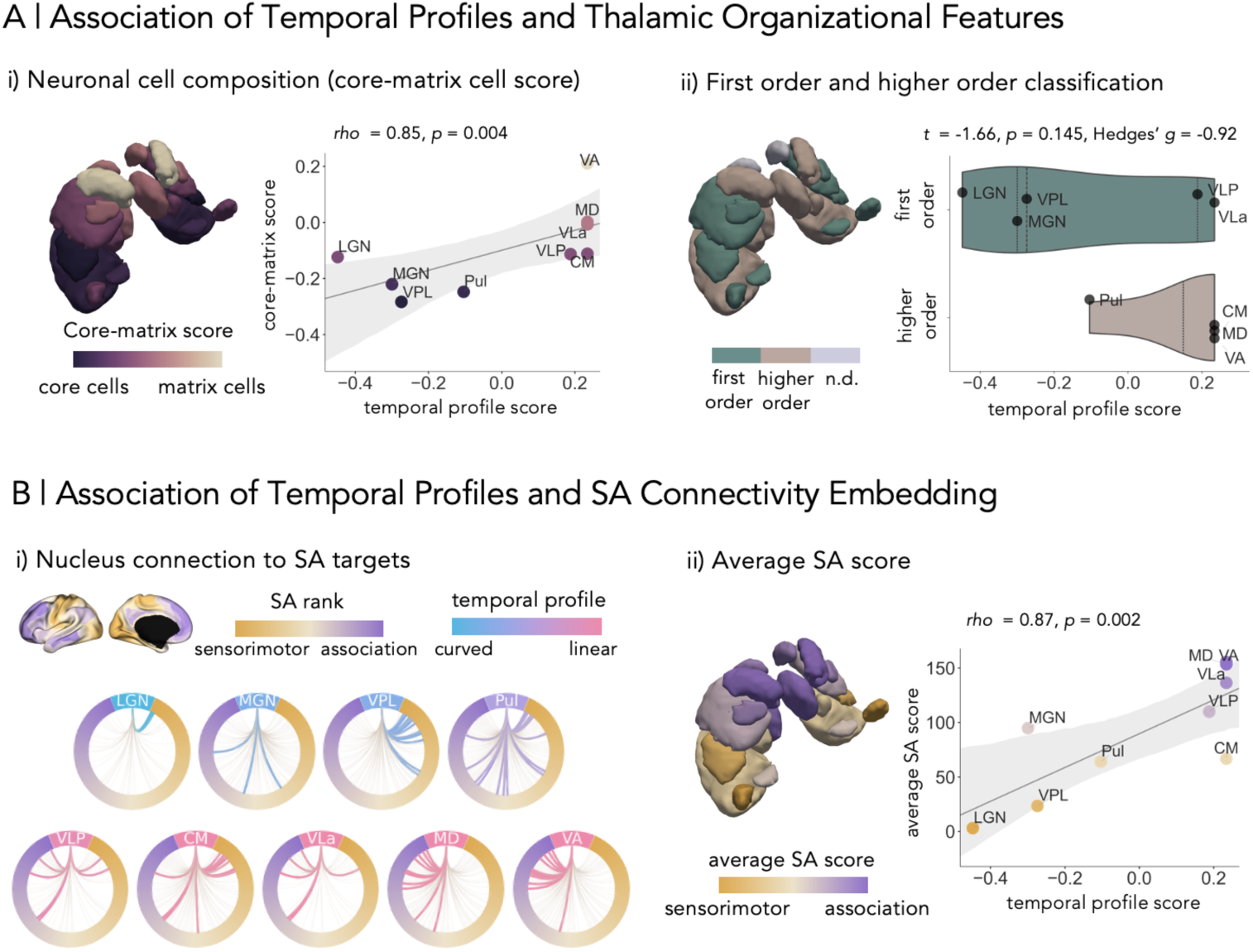
Association between spatiotemporal development of thalamocortical connections and thalamic and cortical features. **A** Temporal profile score (principal component 1 (PC1)) association with thalamic features. i) Core-matrix score (standardized difference between Calbindin and Parvalbumin expression) mapped to thalamic nuclei (left and right hemisphere averaged) (Müller et al., 2020). Scatter plot shows positive relation between temporal profile score and core-matrix score with a linear regression line and a 95% confidence interval. Scatter plot and thalamic nuclei are colored by core-matrix score. Results of Spearman’s correlation and *p*-value are indicated. ii) First order versus higher order classification mapped to nuclei (Sherman, 2005). Relation between temporal profile score and first and higher order classification. Results of *t*-test and Hedge’s *g* are indicated. **B** Association between temporal profile scores and cortical sensorimotor-to-association (SA) axis (Sydnor et al., 2021). i) For each nucleus, the network plot shows the structural connections (colored by temporal profile score) to cortical target regions ordered by SA scores. ii) Per nucleus, the SA average score of nucleus connections is mapped onto the thalamus. Scatter plot shows positive relation between temporal profile scores and SA scores with a linear regression line and a 95% confidence interval. Scatter plot and thalamic nuclei are colored by SA scores. Results of Spearman’s correlation and *p*-value are indicated. Abbreviations: Lateral Geniculate Nucleus (LGN), Medial Geniculate Nucleus (MGN), Ventral Posterior Lateral (VPL), Ventral Lateral Nucleus (posterior) (VLP), Ventral Lateral Nucleus (anterior) (VLa), Pulvinar (Pul), Centromedian Nucleus (CM), Mediodorsal Nucleus (MD), Ventral Anterior Nucleus (VA)

**Fig. 4:**
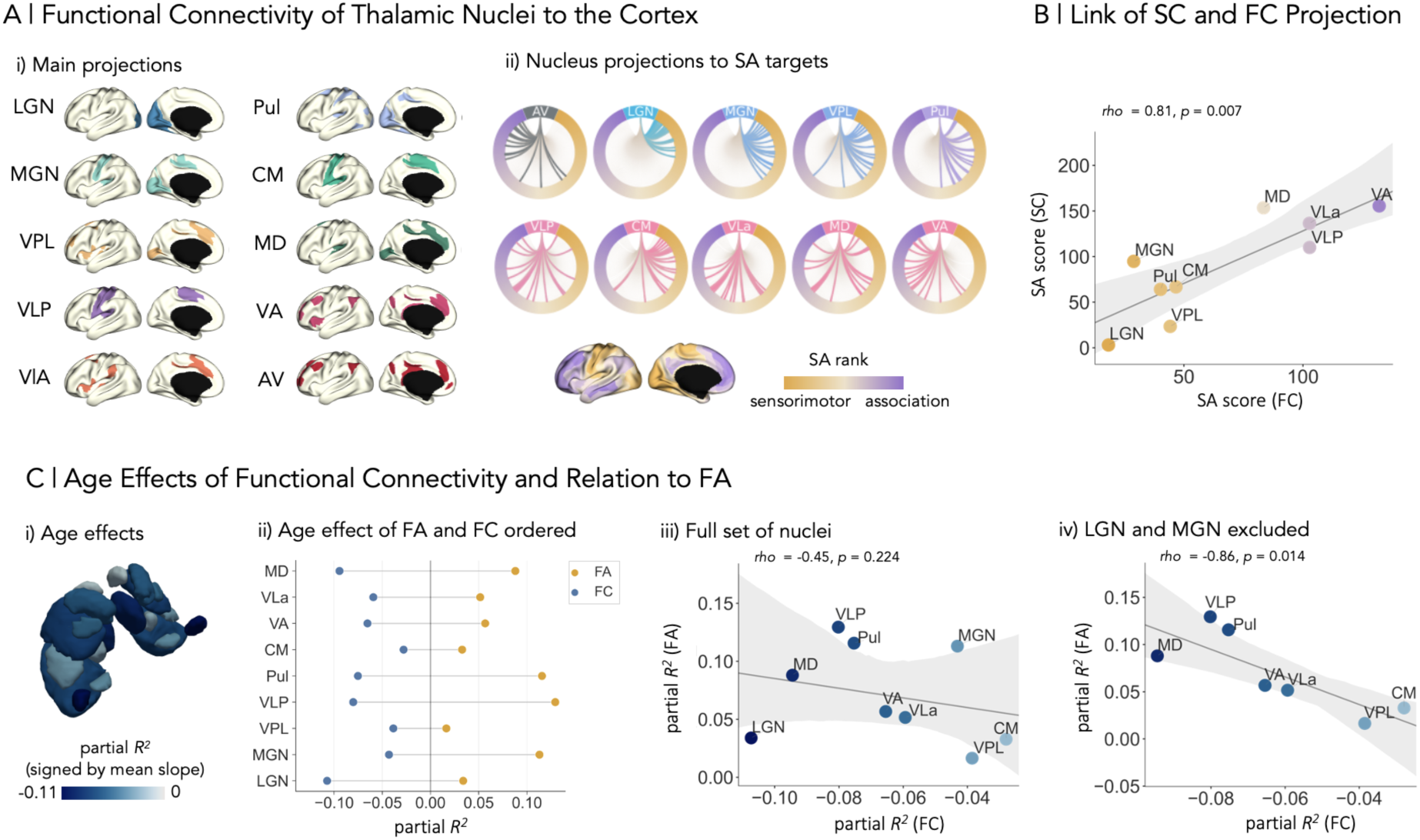
Nucleus-specific thalamocortical functional connectivity maturation. **A** i) Top 10% functional connections for each nucleus mapped onto the cortical surface. ii) Network plots show for each nucleus the functional connected cortical regions ordered by sensorimotor-to-association (SA) scores (Sydnor et al., 2021). **B** Association between SA scores of structural and functional connections. Positive relation is depicted by a linear regression line with a 95% confidence interval. **C** Age effects of functional connections were modeled using generalized additive models (GAMs). i) Partial *R^2^*(signed by mean slope) of each nucleus connection age trajectory are displayed on the thalamus. Comparison of partial *R^2^* (signed by mean slope) of fractional anisotropy (FA) and functional connectivity age trajectories. ii) Nuclei are ordered by normalized absolute difference in partial *R^2^* between FA and FC. iii) Scatter plot shows no relation between nucleus-specific FA-partial *R^2^* and functional connectivity-partial *R^2^* for the full set of nuclei, with a linear regression line and a 95% confidence interval. Scatter is colored by functional connectivity partial *R^2^*. iv) Scatter plot shows relation between nucleus-specific FA-partial *R^2^* and functional connectivity-partial *R^2^* where LGN and MGN being excluded, with a linear regression line and a 95% confidence interval. Scatter is colored by functional connectivity partial *R^2^*. Abbreviations: Lateral Geniculate Nucleus (LGN), Medial Geniculate Nucleus (MGN), Ventral Posterior Lateral (VPL), Ventral Lateral Nucleus (posterior) (VLP), Ventral Lateral Nucleus (anterior) (VLa), Pulvinar (Pul), Centromedian Nucleus (CM), Mediodorsal Nucleus (MD), Ventral Anterior Nucleus (VA), Antero-ventral Nucleus (AV), structural connectivity (SC), functional connectivity (FC)

**Fig. 5:**
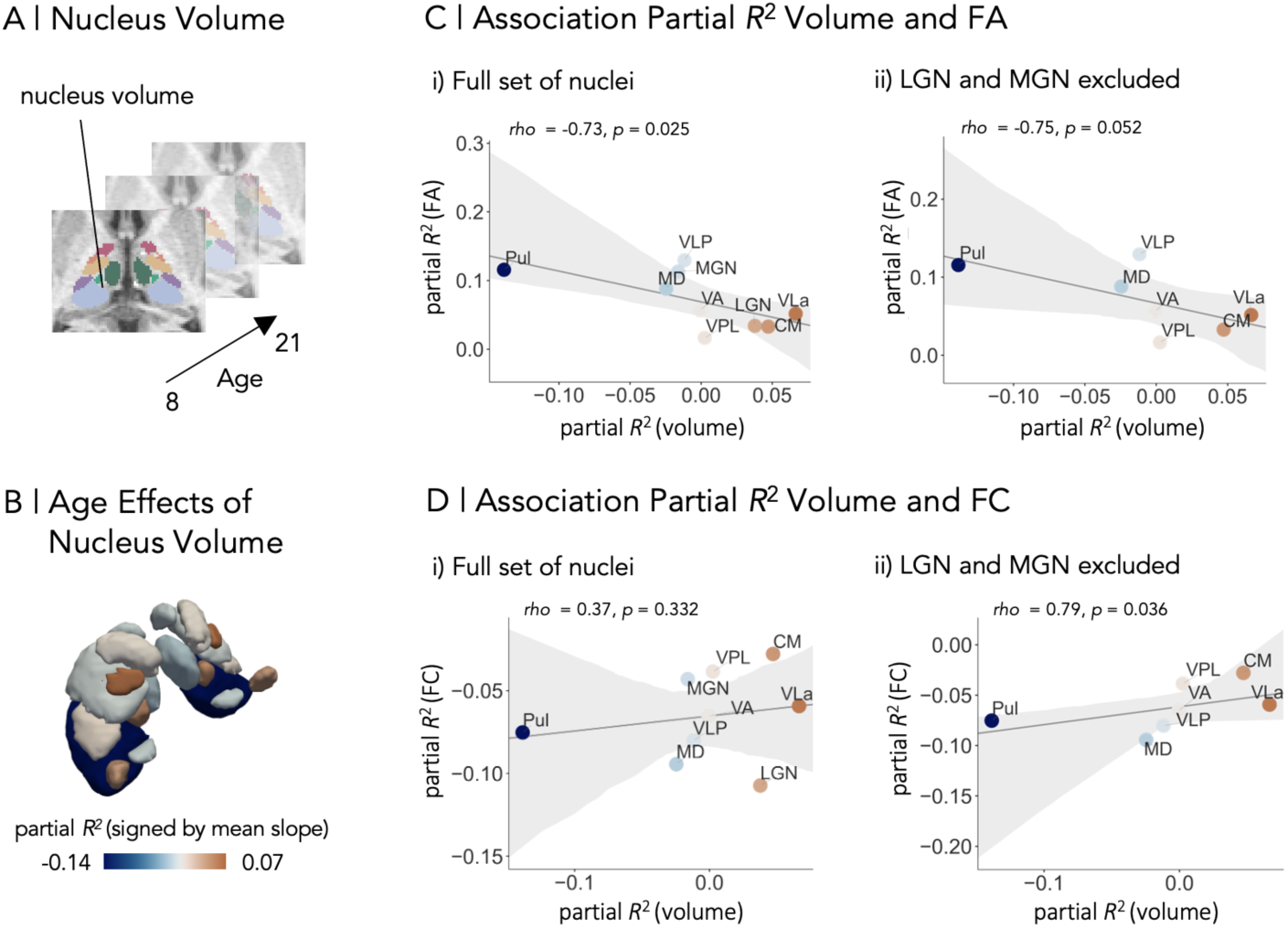
Association between development of nucleus-specific structural and functional thalamocortical connections and nucleus volume maturation. **A** Nucleus volumes were extracted and averaged across hemispheres and their developmental trajectories were modeled using generalized additive models (GAMs). **B** Partial *R^2^,* which describes the magnitude of age effect and is signed by mean slope of the trajectories, was mapped on the thalamus. **C** Relation between volume-partial *R^2^* and fractional anisotropy (FA)-partial *R^2^* of i) the full set of nuclei and ii) the subset with LGN and MGN being excluded, with a linear regression line and a 95% confidence interval. Scatter is colored by volume-partial *R^2^.* D Relation between volume-partial *R^2^* and functional connectivity (FC)-partial *R^2^* of i) the full set of nuclei and ii) the subset with LGN and MGN being excluded, with a linear regression line and a 95% confidence interval. Scatter is colored by volume-partial *R^2^.* Abbreviations: Lateral Geniculate Nucleus (LGN), Medial Geniculate Nucleus (MGN), Ventral Posterior Lateral (VPL), Ventral Lateral Nucleus (posterior) (VLP), Ventral Lateral Nucleus (anterior) (VLa), Pulvinar (Pul), Centromedian Nucleus (CM), Mediodorsal Nucleus (MD), Ventral Anterior Nucleus (VA)

### Age trajectories of whole-thalamus thalamocortical structural connectivity

To establish a robust baseline for our nucleus-specific analysis, we started our investigations by evaluating the developmental trajectories of structural connections between the whole-thalamus and cortical parcels. After filtering spurious connections and applying consensus masking, mean FA was extracted from all connections at the individual level, and averaged across left and right connections to increase robustness. Group-level FA was highly similar across the hemispheres (**Supplementary Fig. 2**).

Age trajectories of the resulting 151 connections were modeled using GAMs, with 83.4% of connections showing a significant age effect, (*p* < 0.05, false discovery rate (FDR)-corrected). The resulting age trajectories were variable in temporal profiles (linear versus curved) and in magnitude of age effects (quantified by partial *R^2^*) across cortical parcels. This variation is illustrated by example connections targeting the visual cortex (V1), a sensory region, and the prefrontal cortex (p47r), an association region (**Fig. 1A**). An overview of the age trajectories of all connections and the spatial distribution of partial *R^2^*, ranging from −0.01 to 0.14, is shown in **Fig. 1A**. To further characterize the trajectories, we calculated their mean first derivatives to quantify the average rate of age-related change. Note that, for interpretability, partial *R^2^* values were assigned a sign reflecting the direction of change (sign of mean slope), such that negative values were referred to an overall decrease of FA with age. Among connections showing a significant age effect, the vast majority exhibited an overall increase of FA within the age range (99.2%), as indicated by a positive mean first derivative. In 11.3% of connections, the modeled age trajectories were significantly non-linear.

In addition to assessing the magnitude of age effects (partial *R^2^*), we sought to characterize variations in timing of development, and thus the shape of the age trajectories. Principal Component Analysis (PCA) provided a data-driven approach to capture nuanced differences in spline shape. Therefore, PCA was applied on the normalized first derivatives sampled along the nucleus-specific trajectories with age-related increase. PC1 explained 79.68% of variance and captured the temporal variations of development (e.g., timing of developmental plateau) by separating trajectories with more linear FA increase from more non-linear trajectories (**Fig. 1B**). Hereafter, PC1 is referred to as the temporal profile component.

Next, we examined the relationships among developmental trajectory related characteristics and their association to the cortical SA axis using Spearman correlations, correcting for spatial autocorrelation using spin permutation (Váša et al., 2018) (**Fig. 1C**). We found that partial *R^2^* closely relates to the mean first derivative of the trajectories (Spearman’s *rho* = 0.96, *p_spin_* < 0.05) but not the temporal profile component (Spearman’s *rho* = −0.16, *p_spin_* = 0.27), suggesting that partial *R^2^* captures the magnitude of age effects but not the temporal variations across trajectories. The temporal profile component, which captures variations of spline shape (e.g., timing of developmental plateau), was related to the SA axis (Spearman’s *rho* = 0.46, *p_spin_* < 0.05), showing that the spatiotemporal development of thalamocortical structural connectivity links to a hierarchical axis of cortical organization as previously reported (Sydnor et al., 2025). Last, to explicitly assess consistency with prior work, we correlated age effects of this study with age effects in (Sydnor et al., 2025), and observed a significant relationship (Spearman’s *rho* = 0.58, *p_spin_* < 0.05) (**Fig. 1D**). These results establish that our tractography pipeline captures known patterns of thalamocortical structural connectivity development at the level of the whole-thalamus, supporting its validity for extending the analysis to the level of individual nuclei.

### Thalamocortical structural connections differ across nuclei and exhibit heterogenous nucleus-specific developmental trajectories

After establishing the developmental trajectories of thalamocortical connections at the whole-thalamus level, we proceeded to examine the developmental trajectories at the level of distinct nuclei. For each individual, the thalamus was parcellated into 10 nuclei per hemisphere using the HIPS-Thomas approach, including: Lateral Geniculate Nucleus (LGN), Medial Geniculate Nucleus (MGN), Ventral Posterior Lateral (VPL), Ventral Lateral Nucleus (posterior) (VLP), Ventral Lateral Nucleus (anterior) (VLa), Pulvinar (Pul), Centromedian Nucleus (CM), Mediodorsal Nucleus (MD), Ventral Anterior Nucleus (VA), Antero-ventral Nucleus (AV) (**Fig. 2A**; (Vidal et al., 2024)).

We first reconstructed the nucleus-specific thalamocortical connections. For each nucleus, we identified their connections with the ipsilateral cortical Glasser parcels (Glasser et al., 2016) using probabilistic tractography. Since nucleus-specific tracking is challenging due to the small size and close proximity of nuclei, we applied a multistage filtering pipeline to ensure nucleus-specific connections and avoid false-positives or signal bleeding from other nuclei, including the removal of spurious streamlines, consensus filtering, and manual exclusion of misdirected streamlines (see methods for details). Next, we generated a combined volumetric tract per nucleus-to-cortex connection. For the majority of nuclei, resulting profiles align with expected anatomical descriptions (see table in (Zhang et al., 2010), summarizing connection profiles from (Jones, 2007)), such as the LGN connections to visual cortical areas, VPL to somatosensory cortical areas, MD and VA connections to prefrontal cortical areas (**Fig. 2B**). Connectivity profiles were highly similar across hemispheres (**Supplementary Fig. 3**).

However, for AV, anticipated connections to limbic and working memory-related regions (Robertson and Kaitz, 1981) were not captured. Instead our results show a connection from AV to orbitofrontal regions. Although we cannot exclude the possibility that this connection, or part of it, reflects a plausible anatomical pathway, the overall reconstruction appeared incomplete, as key components of the AV connectivity were missing. This likely reflects the anatomical location and its surrounding structures that create a challenging tractography environment for streamline propagation. The reconstruction of the thalamocingulate connection has been reported challenging in the past (Weininger et al., 2019), after propagating anterior, streamlines would need to follow a turn, while being biased by the dominant anterior thalamic radiation. Moreover connections through the mammilothalamic bodies to the temporal lobe were not possible to track. Given that AV connection did not sufficiently capture the nucleus-specific expected profile, we report its developmental trajectory descriptively but exclude it from subsequent analysis.

Next, we investigated the unique age-related trajectories of nucleus-specific connections using GAMs. All 10 nuclei connections exhibited significant age effects (*p* < 0.05, FDR-corrected). While across the majority of nucleus connections a general trend of FA age-related increase was observed, the shapes (linear increase versus plateauing) of the trajectories and magnitude of age effects (partial *R^2^*) were nucleus-specific (**Fig. 2C**). A notable exception was observed in the AV connection, which exhibited a decrease of FA followed by an increase, resulting in a u-shaped trajectory (**Fig. 2C**). As reasoned above, the AV profiles are excluded from subsequent analysis.

We next sought to unpack the nucleus-specific developmental trajectories and investigate their maturational patterns from two view points: the temporal variability of nucleus-specific connection development and the magnitude of age effects. To compare the temporal variability of nucleus-specific age trajectories, we performed a PCA on the normalized first derivatives sampled along the trajectories, thereby focusing on spline shape (e.g., timing of developmental plateau) instead of magnitude. PC1, hereafter referred to as temporal profile component, explained 99.45% of variance, and revealed an ordering from trajectories with early increases followed by plateauing (LGN, MGN, VPL, Pul) to trajectories showing approximately linear increases within the age range (VLP, CM, VL, VA, MD) (**Fig. 2D**). For the magnitude of age-related change, nuclei were divided into four groups based on the relation to average rate of change across all nuclei: nuclei connections with greater overall rate of change across the age range (MD, Pul, VLP), early dominant nuclei with greater rate of change in the first half of the age range (MGN, LGN), late dominant nuclei with greater rate of change in the second half of the age range (VA, VLa, CM), and nuclei with lower overall rate of change (VPL) (**Fig. 2E**). The spatial pattern of the magnitude of age effects (partial *R^2^*) mapped onto the nuclei is shown in **Supplementary Fig. 4**. Ultimately, the findings highlight that while an age-related FA increase in cortical connectivity is a common feature of these connections, the temporal patterns of development and magnitude of age effects vary across individual nucleus connections.

### Heterochronous development of nucleus-specific thalamocortical connections is associated with thalamic organization

In the next step, we assessed whether temporal variability in nucleus-specific connectivity development would relate to thalamic features. First, we observed a strong association of the temporal profile component with the underlying distribution of core- and matrix cells within the thalamus (Müller et al., 2020; Hawrylycz et al., 2012), which exhibit different projection patterns (Spearman’s *rho* = 0.85, *p* < 0.05) (**Fig. 3A**). Next, we investigated whether the temporal profile component scores differed between first order and higher order nuclei, which are classified based on the driver source input to the nucleus (Sherman, 2005). Temporal profile component scores were nominally lower in first order (mean −0.12 ± 0.31) compared to higher order nuclei (mean 0.15 ± 0.17), suggesting a tendency toward earlier plateauing in nuclei that receive subcortical driver input and continuous developmental change in nuclei that receive driver input from the cortex, but did not reach statistical significance (Welch’s t-test, *t* = −1.66, *p* = 0.145, Hedges’ *g* = −0.92) (**Fig. 3A**). Given the small number of nuclei in each group, this comparison should be interpreted cautiously.

Last, we investigated how nucleus projections map onto the cortical SA axis (Sydnor et al., 2021), and whether the mean SA score per nucleus is associated with the temporal profile component. As expected, the nuclei varied in their targeting along the SA axis, with some nuclei projecting to a narrow range of regions (*e.g.,* LGN), whereas others exhibited more distributed connections (e.g. MD). To estimate a nucleus-specific SA score, we averaged the targeted SA ranks per nucleus. This mean SA rank showed a significant correlation with the temporal profile component (Spearman’s *rho* = 0.87, *p* < 0.05) (**Fig. 3B**). In contrast, for the magnitude of age effect (partial *R^2^*), we found no association with any of the features (cell distribution (Spearman’s *rho* = −0.05, *p* = 0.898), first order and higher order classification (Welch’s t-test, *t* = −0.15, *p* = 0.88, Hedges’ *g* = −0.09), and SA score (Spearman’s *rho* = 0.37, *p* = 0.332)) (**Supplementary Fig. 5)**. Together, we showed that the relation of temporal features with the SA axis holds at the nucleus level, but can be extended by thalamic cell-distribution and to some extent the first order/higher order classification of the nuclei.

### Nucleus-specific thalamocortical functional connectivity decreases with age and is coupled with structural connectivity age effects in connections with prolonged structural maturation

We next tested whether the maturation in nucleus-specific thalamocortical structural connectivity is related to age-related changes in thalamocortical functional connectivity. Therefore, we first defined nucleus-specific functional connectivity profiles, by calculating the functionally top 10%-strongest connected cortical regions of each nucleus at the group-level (**Fig. 4A**). Connectivity profiles showed high similarity between hemispheres (**Supplementary Fig. 6**). Akin to the structural connections, we mapped per nucleus the functionally strongest connected cortical regions onto the SA axis (**Fig. 4A**), and computed for each nucleus-specific functional connection an average SA score. Structural and functional SA scores were significantly correlated (Spearman’s *rho* = 0.81, *p* < 0.05), indicating a consistent alignment between structural and functional spatial connectivity profiles of each nucleus in the cerebral cortex (**Fig. 4B**). Age effects of functional connections were modeled using GAMs on the cross-hemisphere average of functional connectivity *z*-scores. Thalamocortical functional connectivity for all nuclei showed significant age effects (*p* < 0.05, FDR-corrected) (**Fig. 4C, Supplementary Fig. 7**). While structural connection changes, indexed by FA, showed predominantly increasing trajectories, we found functional connectivity decreasing with age across nuclei (partial *R^2^* signed by mean slope) (**Fig. 4C**). Next, we examined the cross-modality relation between the magnitude of age effects at the nucleus-level. To this end, we ordered the nuclei based on a similarity index, quantified as the normalized absolute difference in partial *R^2^* between modalities. This showed greater correspondence between structural and functional connectivity age effects in nuclei involved in higher order cognition, such as MD and Vla, and more divergent profiles in sensory nuclei connections, such as LGN and MGN (**Fig. 4C**). This was further tested by correlating the magnitude of age effects (partial *R^2^*) between nucleus-specific structural and functional connectivity. Across the full set of nuclei, we found no association between FA and functional connectivity age effects (Spearman’s *rho* = −0.45, *p* = 0.224). However, based on the previous observation, we excluded LGN and MGN, which are both sensory nuclei that decelerate earliest in FA increase. This post-hoc analysis showed a significant correlation (Spearman’s *rho* = −0.86, *p* < 0.05), supporting the pattern of a greater correspondence between structural and functional connectivity in nucleus-connections with prolonged maturation. Temporal profile components between structure and function were not related (Spearman’s *rho* = 0.32, *p* = 0.406) (**Supplementary Fig. 8**). Together, our findings suggest a cross-modality correspondence, not across all nuclei, but of nucleus-connections with prolonged maturation.

### Thalamocortical connectivity maturation relates to nucleus volume development

Finally, we examined how nucleus-specific structural and functional connectivity development was related to changes in age-related change of nucleus volume. To do so, we modeled the age trajectories of nucleus volume using GAMs (**Fig. 5A**). Volumes of 8 nuclei showed a significant age effect (*p* < 0.05, FDR-corrected), whereas volumes of VA and VPL showed no significant age effect (**Supplementary Fig. 9**). We identified overall nucleus volume increase or decrease across the age range by calculating the average slopes along the age trajectories. For the significantly changing nucleus volumes, we found a negative average slope for nucleus volumes of AV, VLP, Pul, MGN and MD, and a positive average slope for nucleus volumes of Vla, LGN, and CM. The whole-thalamus volume decreased significantly with age (*p* < 0.05, FDR-corrected) (**Supplementary Fig. 9**). This suggests that age-related decreases of the whole-thalamus volume masks a heterogeneity at the nucleus level, where some nucleus volumes decrease while others expand. To measure the magnitude of age effects, partial *R^2^* (signed by average slope) was computed (**Fig. 5B**).

Next, we correlated the age effects (partial *R^2^*) of nucleus volume development with age effects (partial *R^2^*) of structural and functional connectivity for the full set of nuclei and the above defined subset of nuclei, with LGN and MGN being excluded. We found age effects of FA and nucleus volumes negatively correlated for the full set of nuclei (Spearman’s *rho* = −0.73, *p* < 0.05), indicating that volume decrease is associated with age-related FA increase (**Fig. 5C**). For the subset of nuclei, this correlation did not reach statistical significance but showed the same trend (Spearman’s *rho* = −0.75, *p* = 0.052) (**Fig. 5C**). For functional connectivity and nucleus volume change, we found no relationship for the full set of nuclei (Spearman’s *rho* = −0.37, *p* = 0.332), for the subset it revealed a relation (Spearman’s *rho* = −0.79, *p* < 0.05). We further applied a PCA on the normalized first derivatives of nucleus volume trajectories to identify temporal patterns, which revealed a PC1 (temporal profile component) with explained variance of 84%. A relation between the temporal profile component of nucleus volume and FA (Spearman’s *rho* = −0.4, *p* = 0.286) and nucleus volume and functional connections (Spearman’s *rho* = −0.67, *p* = 0.05) could not be identified (**Supplementary Fig. 10**). Together, these results indicate that magnitude of age effects, but not the temporal profile of structural and functional connectivity changes, is linked to co-occurring developmental changes in the volume of thalamic nuclei.

## Discussion

In the current work, we mapped and multimodally characterized nucleus-specific thalamocortical structural connectivity maturation *in vivo* across childhood and adolescence. Using probabilistic tractography, we reconstructed the thalamocortical connections of individual nuclei. We found that FA increased across nucleus-specific thalamocortical connections, with differences in the temporal patterns and magnitude of age effects between nuclei. Different nuclei exhibited temporally distinct FA trajectories: nuclei mainly projecting to sensory cortical areas and temporal lobe showed early and rapid increases that plateaued during adolescence (LGN, MGN, VPL, Pulvinar), whereas nuclei mainly projecting to motor, premotor or prefrontal cortical regions showed continued increase throughout development (VLP, CM, Vla, MD, VA). This spatiotemporal pattern was related to thalamic and cortical features, such as cell distribution within the thalamus, the embedding of nucleus-specific connectivity profiles along the SA axis, and nominal differences between first order and higher order nuclei. Notably, the strengthening of structural connections was paralleled by a decrease in functional connectivity across all nuclei, suggesting a decoupling of thalamus and cortex with age. FA and FC age effects were coupled for the subset of nuclei with a more prolonged maturation in structural connections (LGN and MGN excluded), but not for the full set of nuclei. Last, we found the magnitude of structural connectivity age effects being correlated with age-related volume changes in the corresponding thalamic nuclei, suggesting a shared developmental mechanism linking nucleus remodelling and connectivity refinement. The relation was further observed for functional connectivity within the subset of nuclei shown to be coupled to FA age effects. Together, using a multimodal perspective of thalamocortical development, we show different modes of change that are anchored in thalamic organization and nucleus volume development.

White matter pathways between thalamic nuclei and the cerebral cortex support bidirectional communication, enabling bottom-up transmission of sensory and subcortical signals to the cortex and top-down modulation from cortical areas back to the thalamus. Individual thalamic nuclei differ in their connectivity profiles, reflecting their specialized roles in sensory processing, motor control, and higher order cognition (Shine et al., 2023; Antonucci et al., 2021; Zhang et al., 2010; Sherman, 2005; Hwang et al., 2017). Here, we successfully reconstructed nucleus-specific thalamocortical connections, with results showing strong correspondence with known anatomical patterns in the adult brain (see table in (Zhang et al., 2010) derived from (Jones, 2007)). We found LGN connecting to the occipital cortex, MGN connections to the superior temporal lobe, and VPL connections to the postcentral gyrus. VLP, VLa and CM were connected to the motor and premotor cortex. MD and VA targeted the prefrontal cortex. Only AV, a small anterior located nucleus, did not reach anticipated target regions, such as the cingulate (Robertson and Kaitz, 1981), due to methodological limitations related to its location and proximity to other structures. The exact path of the thalamocingulate connection remains debated (Weininger et al., 2019). Likely streamlines that propagated anteriorly failed to sharply turn to the cingulate and proceeded within the dominant anterior thalamic radiation. Moreover, connections from AV via the mamillothalamic bodies could not be reconstructed.

Childhood and adolescence are periods of heightened plasticity, during which the microstructure of perinatally established thalamocortical connections continue to mature (Sydnor et al., 2025; Avery et al., 2022; Alkonyi et al., 2011). FA, which is commonly used to track developmental changes in structural connections, serves as a measure of white matter microstructure (Lebel and Beaulieu, 2011; Lebel et al., 2008; Conte et al., 2024; Reynolds et al., 2019; Barnea-Goraly et al., 2005) and is sensitive to different biophysical aspects, such as myelin content, the axon diameter, cell density, membrane permeability, and water content (Tamnes et al., 2017; Beaulieu, 2002). Although FA is not specific to a single underlying biological process, developmental changes of FA are commonly attributed to axon myelination and growth in axon diameter, which makes neural transmission more efficient (Paus, 2010). Therefore, the overall increase of FA suggests a strengthening or refinement of the thalamocortical structural connections.

Importantly, the nucleus-specific thalamocortical connections showed heterogeneous developmental trajectories, varying in both the temporal patterns of maturation and the magnitude of FA-age effects. In particular, the variance of temporal patterns across nucleus-specific connection development aligned with organizational principles of the thalamus, including core- and matrix cell distribution and projection patterns to the cortical SA axis. Thalamocortical connections of thalamic nuclei that are involved in sensory functions and project to sensory cortical areas (LGN-visual, MGN-auditory, VPL-somatosensory) showed an increase of FA in childhood and early adolescence that stabilized in late adolescence. These nuclei tend to have a relatively higher proportion of core cells, which project targeted to the cortex, suggesting that white matter connections including axons of core cells mature in childhood and adolescence but stabilize in late adolescence. How much the connections changed within the childhood and early adolescence varied between nuclei: MGN and LGN connections displayed the most pronounced changes during childhood and early adolescence, whereas VPL connections showed the least change throughout the whole age range. This demonstrates that connections of sensory nuclei change early in development but vary in rate of change. The Pulvinar connections showed a later deceleration and overall a high increase across the age range. While this nucleus, similar to the sensory nuclei, exhibits a higher proportion of core cells, the Pulvinar is connected to visual cortex areas, but also cortical association areas, and involved in cognitive functions, such as visual attention (Zhou et al., 2016). By contrast, connections of nuclei that project to motor- and premotor (VLP, Vla, CM), and prefrontal cortex (MD,VA) exhibit relatively linearly increasing age trajectories, with VLP and MD showing the highest change. The high proportion of diffusely projecting matrix cells in nuclei VA, Vla and MD may help to explain the prolonged maturation of their connections, as matrix cell projections preferentially target cortical layer I-III and are more involved in higher cognitive processes (Müller et al., 2023, 2020). Indeed, a proceeding maturation of thalamocortical connections within this age range is in line with reported changes of white matter that showed maturation until the third decade of life (Lebel and Beaulieu, 2011). Overall, these results support the hypothesis that the timing of thalamocortical maturation reflects functional demands of their cortical projection targets in the cortex, and relate to aspects of intrinsic thalamic organization, such as neuronal cell types. Of particular note, we found higher rates of change in the connections of MD, a nucleus involved in functions such as cognitive control and executive function (Wolff and Halassa, 2024; Ferguson and Delevich, 2020) that get particularly refined in adolescence.

The temporal development of nucleus-resolved tracts was aligned with the spatiotemporal pattern of cortical development along the SA axis (Sydnor et al., 2021), where hierarchical timing has been observed, for example, in cortical myelination, cortical thinning and surface expansion (Baum et al., 2020; Grydeland et al., 2019; Flechsig, 1901; Sotiras et al., 2017; Hill et al., 2010; Sydnor et al., 2021), and has also been reported for cortico-cortical structural connectivity development (Xu et al., 2026). Our findings are in line with previous work (Sydnor et al., 2025), showing that temporal patterns of developmental trajectories of the whole-thalamus connections align with their target area within the SA axis. Connections that target areas in the sensory cortex reach a plateau of FA increases earlier than connections to association areas. Our findings extend this framework at the nucleus level; thalamocortical connections targeting from sensory nuclei mature earlier, while nuclei projecting to association areas show prolonged development, reinforcing the notion that thalamocortical maturation is not isolated but coordinated with cortical development.

Additionally, we considered the classical distinction into first- and higher-order thalamic nuclei (Sherman, 2005). First order nuclei receive their driver input from the subcortex and relay sensory input to the cortex, while higher order nuclei receive ‘driver’ input from the cortex, and are more involved in integrative and cognitive functions. In rats, the higher order circuits show elevated expression of GAP-43, a protein associated with growth and neuronal plasticity, suggesting greater developmental malleability (Feig, 2005, 2004). While we observed a trend toward early consolidation in first order nuclei, and more sustained development in higher order nuclei, this difference did not reach significance and should be interpreted cautiously due to the small sample size within the groups (first order: 5 nuclei, higher order: 4 nuclei). Moreover, the binary classification might not fully capture across-nucleus variability in connectivity development, particularly compared with more continuous measures of thalamic and cortical organization.

Structure and function are in continuous interplay, with structure constraining functional dynamics and functional activity contributing to structural changes (Honey et al., 2007; Draganski et al., 2004). Therefore, functional connectivity offers a complementary perspective to structural connectivity development, providing additional insights on thalamocortical maturation. Overall, structural and functional nucleus-specific connectivity profiles were organized in a strongly similar manner along the SA axis, indicating a shared large-scale organizational principle. Deviations between the connectivity profiles across modalities reflect modality-specific sensitivities. For example, tractography revealed connections between VPL and somatosensory cortex, whereas this was absent in functional connectivity, consistent with a previous study (Zhang et al., 2010). With respect to maturation, we found a decrease in functional connectivity across all nuclei with age, revealing that thalamocortical interactions undergo substantial age-related changes. The observed pattern is in line with previous work reporting thalamocortical or subcortical-cortical decrease in functional connectivity in development (Badke D’Andrea et al., 2023; van Duijvenvoorde et al., 2019; Huang et al., 2021; Supekar et al., 2009). However, it contrasts studies that show heterogeneous network-specific age-related patterns of functional connectivity, with a decrease in some networks but increase in others (Váša et al., 2020; Fair et al., 2010; Steiner et al., 2020). Our finding suggests that the strengthening of structural connections with age is paralleled by a reduced baseline coupling between thalamus and cortex with age. Age effects of FA and functional connectivity were correlated in nuclei with a more prolonged maturation of structural connections, which suggest a co-maturation in those nucleus connections. This association was attenuated when including LGN and MGN, as these nuclei showed strong age effect in one modality but weak effects in the other. A possible interpretation for the decrease of thalamocortical connectivity is that the role of thalamocortical interaction shifts within development from an intrinsical organizational to a selective context-dependent role. While in early development spontaneous thalamic activity contributes to cortical development (Antón-Bolaños et al., 2018), in childhood and adolescence the thalamus de-engages at rest which might reflect the contribution to network segregation and specificity (Park et al., 2024). This might align with the paralleled increase in transmission efficiency in structural connections. Together, our findings underline the importance of a multimodal approach to understand the role of thalamocortical interplay in development.

Finally, we sought to investigate the hypothesis that maturation of structural and functional thalamocortical connectivity is anchored in developmental processes within the corresponding nuclei. Developmental changes of thalamic nuclei have previously been reported using volumetric MRI (Huang et al., 2024; Raznahan et al., 2014; Thalhammer et al., 2024a), and abnormalities in nucleus volume have been linked to neurodevelopmental disorders, such as schizophrenia (Huang et al., 2024, 2020; Elvira et al., 2025). This proposes volumetric change as a macrostructural measure for nucleus maturation. Although volumetric changes cannot be directly linked to underlying biological maturational processes, they may relate to microstructural reorganisation, such as dendritic pruning or a reduction of neuronal density, and connectivity. Consistent with prior work (Carrick et al., 2025; Sussman et al., 2016; Huang et al., 2024; Herting et al., 2018), we found decreases in the whole-thalamus volume with age. However, age trajectories of individual nuclei differed, with some nuclei showing contraction, while others showed expansion or stability (Huang et al., 2024; Raznahan et al., 2014), underscoring the importance of a nucleus-resolved approach. Previous studies reported a volume reduction for VL, VA and MD, similar to our findings, and proposed expansion and reduction of nuclei to be circuit-specific (Raznahan et al., 2014). Crucially, we found that a reduction in nucleus volume was associated with a greater age effect in FA in its corresponding connection and for the subset of nuclei (with LGN and MGN being excluded) more decrease in functional connectivity. Conversely, nuclei that expanded in volume showed smaller FA age effects and for the subset of nuclei (with LGN and MGN being excluded) less decrease in functional connectivity. One possible interpretation is that volume decrease may reflect a reduction of neurons and synaptic pruning of redundant circuitry, leading to a more efficient selective connectivity (as seen by increase in FA). Postmortem work in humans underlines this interpretation of nucleus reorganization by reporting a decrease of neurons (41%) in MD in adults compared to newborns (Abitz et al., 2007). Moreover, in the rats ventral medial prefrontal cortex a loss of neurons has been shown together with a reduction in volume during adolescence, supporting a link between volume change and neuron density (Markham et al., 2007). From a behavioral perspective, an association between volume reduction (in MD and Pulvinar) and increasing scores in executive function have been reported (Huang et al., 2024), suggesting that microstructural reorganisation might be related to cognitive refinement. Therefore, remodelling mechanisms within the thalamic nuclei may anchor structural connectivity refinement. More speculatively, a reduction of neurons in the nuclei could reflect maturational specialization, potentially contributing to lower functional connectivity through increasing network segregation. Future work at the histological microscale is needed to understand the underlying biological processes and to support causal inference. It should be noted that nuclei are not only connected to the cortex, but are embedded in broader subcortical networks including connections with the basal ganglia and the cerebellum, which through their interaction may also have influence on nucleus development. However, taken together, the association between nucleus-level volume change and changes in the corresponding thalamocortical connections suggests an underlying mechanistic link in the co-maturation of nuclei and their connections.

While our results provide novel insights into thalamocortical development, several limitations must be acknowledged. Our study used a cross-sectional design of HCP-D which limits inferences about within-individual developmental trajectories. The dataset offers a large sample size in combination with high quality diffusion MRI data, but longitudinal data would be needed in future work to fully capture within-person change. Further, a crucial step in our study was the precise construction of nucleus-specific connections using tractography, which enables an indirect estimate of white matter pathways *in vivo* but is not able to resolve axonal connections at the microscopic level. Hence, tractography cannot account for directionality of the approximated connection, and the term ‘target region’ used in this study describes how connections are defined methodologically rather than a biological direction. In fact, animal research has shown that connections from the cortex to the thalamus outnumber the ones from thalamus to cortex, highlighting the relevance of feedback loops (Bagshaw, 2026; Mumford, 1992; Briggs, 2020; Sherman and Koch, 1986). In addition, tractography has well-known limitations, such as resolving ‘crossing-fibers’ and a tendency to generate false-positive connections. We mitigated these issues by employing probabilistic tractography based on the ball-and sticks model with 3 possible fiber directions and strict filtering of potential false-positives. Nevertheless, inaccuracies can not be excluded fully using tractography. Further, due to the size, location, and proximity of nuclei, signal leakage is another concern in both diffusion and functional MRI. The reconstruction of the thalamocingulate connection expected from AV was not successful, likely due the required sharp turn of the propagating streamlines into the cingulum combined with a strong overshadowing signal of the dominant anterior thalamic radiation (see review (Weininger et al., 2019) for details). Microstructural properties of the defined connections were measured using FA, which reflects diffusion directionality within a voxel but lacks fiber specificity, meaning that portions of the FA values may be influenced by intersecting pathways in the same voxels. Complementary approaches, such as invasive animal research are essential to resolve thalamocortical development at the cellular level.

In conclusion, by resolving thalamocortical structural connections at the level of individual nuclei, this study highlights thalamocortical development as a nucleus-specific process and enabled us to demonstrate that variations of maturational pattern of nucleus-specific thalamocortical connections are linked to organizational, developmental and functional heterogeneity of thalamic nuclei. This offers a framework relevant to understanding nucleus-specific contributions to neurocognitive development in childhood and adolescence. Further longitudinal studies will be essential to track individual trajectories and link them to cognitive development and behavioral outcomes in healthy and clinical populations.

## Methods

### Dataset HCP-D

#### Sample

For this study, we used the cross-sectional developmental sample of the Human Connectome Project in Development (HCP-D, release 2.0, (Somerville et al., 2018)) that was downloaded from the National Institute of Mental Health Data Archive (NDA). HCP-D aimed to recruit a sex-balanced sample matching the diversity in US-population by ethnicity, race, and socioeconomic status. Substantial exclusion criteria for participation were insufficient English fluency, MRI contraindications, premature birth, serious medical, endocrine, or neurological conditions, hospitalization > 2 days for certain physical or psychiatric conditions or substance use and treatment > 12 months for psychiatric conditions (detailed exclusion criteria are described in (Somerville et al., 2018). Data was acquired at 4 sites in the USA (Boston, Los Angeles, Minneapolis, St. Louis) with 3T Siemens Prisma scanners and matching scanning protocols. Informed written consent for participation was obtained from participants or from parents or legal guardians, in case participants were younger than 18.

In this study, we included data from 604 participants aged 8-21 years (mean age of 14.7 ± 3.9 years, 323 female, 281 male, **Supplementary Fig 1**). Children between 5 and 8 years were excluded due to the small sample size and differing image acquisition parameters in this age range. To ensure high data quality, the sample was restricted to participants with complete diffusion weighted imaging runs, and further exclusions based on preprocessing quality control criteria (see Preprocessing section).

#### Image acquisition

Acquisition details are described in detail elsewhere (Harms et al., 2018), but in short include the following: T1 Weighted (T1W) images were acquired using a 3D multi-echo MPRAGE sequence with the following parameters: TR = 2,500 ms, TI: 1,000 ms, TE = 1.8/3.6/5.4/7.2 ms, flip angle: 8 degrees, 208 slices, voxel resolution: 0.8 mm isotropic. Diffusion weighted images (DWI) were acquired in four consecutive runs with a multiband factor of 4, and 2 shells of b = 1,500 and 3,000 s/mm^2^ in 185 distinct diffusion-weighting directions. Each direction was acquired twice with opposite phase encoding directions (AP and PA) to enable robust distortion correction. 28 b = 0 s/mm^2^ images were acquired equally interspersed across the 4 runs. Imaging parameters included a TR of 3,230 ms and an isotropic voxel resolution of 1.5 mm. Resting-state fMRI (rs-fMRI) data was acquired in four runs using a 2D multiband (MB) gradient-recalled echo echo-planar imaging sequence with the following parameters: MB8, TR = 800 ms, TE = 37 ms, flip angle: 52 degree, voxel resolution: 2 mm isotropic. Runs were acquired for 26 mins (total) in opposite phase encoding (AP and PA) direction. Participants were instructed to stay awake and focus on a fixation cross.

### Preprocessing

Preprocessed T1W and rs-fMRI data were provided by HCP-D, generated by following the HCP Pipelines (v. 3.22). As the full preprocessing procedures are described elsewhere (Glasser et al., 2013; Van Essen et al., 2013; Smith et al., 2013), only a brief summary is provided here. In contrast, DWI was processed using our custom pipeline, described in detail below.

#### Structural image preprocessing

Briefly, T1W images underwent a gradient distortion correction, bias-field correction and registration to MNI space. This was followed by processing the native space images using the HCP-FreeSurfer pipeline, which is based on FreeSurfers’s (v. 5.2) *recon-all* workflow but incorporates adaptations for the high-resolution HCP data, including improved brain extraction and final white and pial surface placement using the native high-resolution images. The pipeline produces anatomical segmentations, reconstruction of cortical surfaces and surface-based registration. In a final step, the surfaces were converted to HCP-standard space and cross-subject cortical alignment was refined using multimodal surface matching (Glasser et al., 2013; Van Essen et al., 2013).

#### DWI preprocessing

For preprocessing the DWI data following our custom pipeline, we used built-in functions of FSL (v. 6.0.3 and 6.0.7.7 (Jenkinson et al., 2012)) and MRTrix (v. 3.0.3 (Tournier et al., 2019)). After denoising the data (MRTrix dwidenoise, (Veraart et al., 2016a, 2016b; Cordero-Grande et al., 2019), Gibbs ringing artefacts were removed (MRTrix mrdegibbs, (Kellner et al., 2016). The susceptibility-induced off-resonance distortion fields measured using the two reversed phase-encoding directions (AP and PA) were estimated using FSL topup on the gibbs unringed single-band EPI reference scans (FSL v. 6.0.3, (Andersson et al., 2003; Smith et al., 2004)). For the following step, a brainmask was created using FSL BET (v. 6.0.3) and visually quality controlled. To correct for eddy current-induced distortions, subject movement, and replacement of slices that were detected as outliers, we run eddy_cuda10.2 (here FSL v. 6.0.7.7 was used due to compatibility with cuda10.2 installed at the GPUs, (Andersson and Sotiropoulos, 2016; Andersson et al., 2016)) using the off-resonance field estimated by topup as an input. The in-built FSL eddy quality control was performed for quality checks on the subject- and grouplevel (v. 6.0.3 eddy_quad and eddy_squad (Bastiani et al., 2019)). Quality control led to the exclusion of participants with relative motion exceeding 4 standard deviations (0.66 mm), and one additional participant who failed visual inspection due to a large percentage of outliers in the b = 1,500 s/mm² volumes (4.23%).

#### rs-fMRI preprocessing

In brief, the rs-fMRI data underwent correction for gradient nonlinearity induced distortion, motion correction using the single-band reference image (6 degrees of freedom), EPI distortion correction using topup, registration to T1W and non-linear registration to MNI 152 space, intensity normalization and bias field removal. The data was then mapped to the standard CIFTI gray-ordinate space (stores cortical surface as vertices and subcortex as volume voxel) and smoothed with a 2-mm FWHM Gaussian kernel (Glasser et al., 2013; Van Essen et al., 2013; Smith et al., 2013). We decided against additional stronger smoothing to limit partial-volume effects between neighbouring thalamic nuclei. On top of the HCP minimally preprocessing pipeline, we applied bandpass filtering (0.01 - 0.1 Hz) and detrending using the Python package *nilearn* (v. 0.11.0). One participant was excluded due to excessive head motion, defined by relative root-mean-square (RMS) displacement > 0.25 mm (threshold defined by > 4 SD above the group-mean), resulting in 603 participants for our functional analysis.

### Thalamus segmentation

Thalamic nuclei were segmented using the HIPS-THOMAS algorithm (Vidal et al., 2024), which provides automated, subject-specific thalamic parcellation based on T1W images. HIPS-THOMAS overcomes poor intrathalamic contrast in standard T1W images by synthesizing a white-matter-nulled like image contrast from T1W images using histogram-based polynomial synthesis (HIPS) yielding in 12 subregions close to the Morel atlas (Williams et al., 2024; Krauth et al., 2010). This approach provides the optimal balance between state-of-the-art segmentation accuracy of nuclei segmentation (Williams et al., 2024), anatomical detail and practicality such as the number and size of nuclei in regard to our resolution. Beyond that, HIPS-THOMAS has previously been successfully applied across a broad age range spanning from childhood to late adulthood in normative and clinical cohorts, supporting its applicability across the age range in our study (Young et al., 2025; Williams et al., 2024). As input we used the non-intensity normalized T1W image in native freesurfer space. The segmentation yielded 10 thalamic nuclei: Antero-ventral Nucleus (AV), Ventral Anterior Nucleus (VA), Ventral Lateral Nucleus (anterior) (VLa), Ventral Lateral Nucleus (posterior) (VLP), Ventral Posterior Lateral (VPL), Pulvinar (Pul), Lateral Geniculate Nucleus (LGN), Medial Geniculate Nucleus (MGN), Centromedian Nucleus (CM), and Mediodorsal Nucleus (MD). Structures not corresponding to thalamic nuclei were excluded (Mammillothalamic Tract and Habenula). For quality control, segmentations with particular small or large thalamus volumes relative to the average were visually inspected.

### Structural connectivity between thalamic nuclei and cortex parcels

#### Thalamic seed and cortical target mask

As seed masks for the tracking of structural connections we used the above described segmentation of 10 thalamic nuclei in native freesurfer space. Cortical target regions were defined using the multimodal Glasser parcellation, including 180 regions per hemisphere and which was built on multiple imaging modalities (Glasser et al., 2016). For tractography, cortical target masks in voxelspace were generated in native freesurfer space (matching the thalamus segmentation) and slightly shifted into the white matter to facilitate streamline termination. We used Freesurfer (v. 7.4.1, (Fischl, 2012)) to first project the Glasser annotation in fsaverage5 onto the white matter/grey matter boundary in native freesurfer T1W space that was shifted 0.5 mm into the white matter.

#### Tractography

Tracking structural connections between thalamic nuclei and cortical parcels is challenging due to complex fiber architecture, the dense packing of adjacent nuclei, and the deep anatomical location of the thalamus. To address these challenges and enable nuclei-specific thalamocortical connectivity reconstruction, we apply probabilistic tractography, which models uncertainty of fiber orientations in a voxel and samples streamline trajectories probabilistically, improving sensibility to plausible pathways (Behrens et al., 2007, 2003). To estimate the distribution of diffusion directions while accounting for crossing fibers in each voxel, we ran FSL’s *bedpostx_gpu* (FSL v. 6.0.7.7, cuda v. 12.0) with a default of 3 fiber directions and using the multi-shell extension of the ball- and sticks model (Jbabdi et al., 2012).

Next, probabilistic fiber tracking between thalamic nuclei and ipsilateral cortical parcels was computed using FSL’s *probtrackx2_gpu10.2* (FSL v. 6.0.7.12, cuda v. 11.2). Tracking was initialized in each thalamic nucleus with 20000 samples per voxel (step length = 0.5 mm, curvature threshold = 0.2, number of steps = 2000, loopcheck, oneway condition). Thalamic seed- and cortical target masks were provided in native freesurfer space. To this end, co-registration between native freesurfer space and diffusion space was computed using FSL *FLIRT* (v. 6.0.3) with 6 degrees of freedom and mutual info as the cost function. The transformation was provided to *probtrackx2_gpu10.2*. Tractography was constrained using a tracking mask that encompassed the thalamus and white matter (dilated by 1mm), while excluding the corpus callosum to ensure intra-hemispheric tracking, the ventricles, and the cerebellum. Probtrackx produces two complementary outputs for each nuclei-cortical parcel combination in each hemisphere: (i) number of streamline counts that reach the target parcel, which we used to define nuclei-specific connections and (ii) a tract density map (fdt_paths.nii.gz) representing the number of streamlines passing through each voxel and reaching the target, which we used to reconstruct the volumetric connections.

#### Construction of nucleus-specific thalamocortical structural connectomes

For defining the nucleus-specific cortical target parcels, we used the number of streamline counts reaching each target. Therefore, we extracted the streamline counts for each thalamic nucleus-to-cortical parcel combination at the participant level. To avoid a bias due to changes of nucleus size with age, we calculated the relative streamline count by dividing absolute streamline count by nucleus volume. To ensure analyzing anatomically meaningful and robust connections, we applied systematic filtering steps at different stages in our pipeline. Since probabilistic tractography is prone to generate spurious connections driven by noise and model uncertainty, we removed spurious connections under a certain threshold, tailored to each nucleus (nucleus-cortical parcel connections below 10 streamlines for AV, VA, VPL, Pul, MGN, CM, MD and 40 streamlines for Vla, VLP and LGN). The remaining relative streamline counts were log-transformed to account for their skewed distribution. Robustness was further enhanced by applying a consistency-based thresholding, retaining only connections present in at least 90% of participants (**Supplementary Fig. 11**).

The dense packing of thalamic nuclei poses a risk of non-specific connection arising from signal bleeding from neighbouring nuclei tracts. Given the study’s aim of analysing nucleus-specific connections, it was important to minimize this risk by careful selection of connections. To this end, we used a two-step procedure including thresholding followed by anatomically informed refinement for some nuclei: After computing a group-average connectivity matrix by averaging the filtered log-transformed relative streamline counts, for each nucleus we selected the top 10% strongest connections using percentile-based thresholding in the right and left hemisphere separately. For the MGN this yielded the expected connection targeting the superior temporal gyrus, but also revealed overlap with LGN patterns likely due to signal bleeding because of their close spatial proximity. Therefore, we removed the LGN connections that appeared in the MGN mask. From the LGN profile we excluded the connection to the temporal pole. This connection likely arose from the sharp curve of the Meyers loops in the anterior temporal lobe (part of the optic radiation, (Chamberland et al., 2018) where streamlines mistakenly propagate to the temporal pole. Lastly, we excluded the temporal lobe connection of AV, VA and MD. A careful examination of the reconstructed pathway revealed a partial volume effect that leads to false-positive connections of those nuclei via the adjacent white matter of the fornix and the anterior commissure to the temporal lobe. (**Supplementary Fig. 11**).

#### Generation of whole-thalamus and nucleus-specific volumetric tracts

After defining the nucleus-specific cortical target parcels, we used the tract density maps to generate volumetric tracts for each participant for each nucleus-to-parcel combination. Low-density voxels with fewer than 10 streamlines were considered as noise and removed. From the remaining voxels the core of the tract is then defined by retaining voxels within the top 10% of streamline density. A binary mask of the core tract is generated excluding the thalamus.

For each nucleus and hemisphere, the binary masks of the before defined connected parcels are merged to one volumetric tract. For ‘whole-thalamus’ analysis, connections were considered valid if they survived consistency thresholding in at least one nucleus. For each parcel, we then combine the above defined volumetric tracts of all nuclei that showed a connection after consistency thresholding.

#### Fractional Anisotropy (FA)

For investigating age-related changes of the whole-thalamus and nucleus-specific structural connections, we used FA. FA is a reliable microstructural measure that quantifies the degree of diffusion anisotropy. The metric is well suited to capture white matter age effects and has been used to study maturation of structural connections (Lebel and Beaulieu, 2011; Lebel and Deoni, 2018) and specifically whole-thalamus thalamocortical connections (Sydnor et al., 2025). To obtain FA maps for each participant, we first fitted diffusion tensors to each voxel using FSL’s *dtifit* (v. 6.0.3) with the weighted least squares option that accounts better for heteroscedastic noise. For quality control, we visually inspected the FA maps and sum of squared error (SSE) maps that reflect the goodness of fit of the tensor model in each voxel. FA maps were then aligned to native freesurfer T1W space using FSL *FLIRT* with mutual information cost function and trilinear interpolation to match the volumetric tracts. For each participant, we then extracted the average FA within the whole-thalamus and nuclei-specific volumetric tract. For modeling age effects, we then averaged the corresponding FA from the right and left hemisphere.

### Functional connectivity between thalamic nuclei and cortex parcels

#### Thalamic seed and cortical target mask

Functional connectivity was computed based on the MSMAll registered Cifti files (dtseries.nii) in MNI nonlinear space. Therefore, we warped the thalamic segmentation from native freesurfer T1W space to the MNI 152 nonlinear 2 mm space using the HCP-provided transformation files. To enhance robustness, we generated a consensus thalamus mask aggregating the warped subject-specific segmentations and retained only the voxels with cross-subject agreement defined by higher than the 70th percentile (voxels with high consistent overlap between subjects). Time series of BOLD activity was then averaged across nuclei voxels and for whole-thalamus analysis across all thalamic voxels. Parcellating the cortical functional data into 360 Glasser (Glasser et al., 2016) parcels was done using the Python package *hcp-utils* (v. 0.1.0)

#### Functional connectivity computation

For each hemisphere and each subject, functional connectivity was calculated as the Pearson correlation of the BOLD time series between pairs of thalamic nuclei and the ipsilateral cortical parcels. Correlation coefficients were then Fisher z-transformed. A group-level functional connectivity matrix was generated by averaging the nucleus-to-cortex functional connectivity matrices across all subjects. For each nucleus and for each hemisphere separately, the top 10% connected parcels were identified using the 90^th^ percentile of connectivity values as a threshold. Based on this threshold, a binary mask was created for each nucleus, indicating the nucleus-specific strongest connections. For each subject and for hemisphere, the nucleus-specific functional connectivity of the identified parcels was averaged. Finally, left and right hemisphere nucleus-to-cortex functional connectivity values were averaged for modeling.

### Nucleus volume

The volume of segmented thalamic nuclei and the whole-thalamus was output of the HIPS-THOMAS approach (Vidal et al., 2024). For model input, we average the nuclei and whole-thalamus volumes of the left and right hemispheres.

### Modeling of age effects using generalized additive models

Age-related effects of structural thalamocortical connections, functional thalamocortical connections, and nucleus volumes, were assessed using GAMs. We therefore used the *mgcv* (v. 1.9-1) and *gratia* package (v. 0.10.0) in R (v. 4.5.0) (Wood, 2017; Simpson, 2024). GAMs offer a high flexibility in capturing linear and nonlinear relationships and are therefore widely used and well suited for characterizing developmental trajectories. For structural connectivity, functional connectivity, and nucleus volume, we fit GAMs with the respective metric (FA, functional connectivity correlation coefficient, nucleus volume) as the dependent variable, age as smooth term, and sex and motion (for FA, functional connectivity) as linear covariates. To estimate the smooth function, thin plate regression splines were used, with smoothing parameters estimated based on a restricted maximum likelihood (REML). Each GAM estimates a smooth function for age as a weighted combination of basis functions. The parameter *k* defines the basis dimensions, and thus limits the degrees of freedom (*k*-1) for the smooth function. Put differently, it constrains the flexibility of the curve. Given the modest complexity of our data, we used a penalized (*fx* = False) spline basis of *k* = 3, which was sufficient to capture the patterns without overfitting. The choice of *k* = 3 is consistent with prior literature (Sydnor et al., 2025). For each model fit, we exclude subjects where the response variable differs more than 3 standard deviations from the mean (between 0 and 10 subjects excluded). Significance of the smooth age term was assessed using F-tests implemented in *mgcv*, with corresponding *p*-values indicating whether the smooth term significantly improves model fit. *P*-values were FDR-corrected using the Benjamin-Hochberg procedure. Further, we formally tested whether developmental trajectories were significantly non-linear. This was performed by fitting GAMs that included both a linear and a smooth age term constrained to capture only nonlinear deviations. A non-significant *p*-value of the smooth term indicated that a linear model described the age effect adequately.

### Temporal profiles and magnitude of age effects

Partial *R^2^* was calculated to describe the contribution of age to the models. This was approached by fitting a covariate-only null model and computing the proportion of variance explained by age from the reduction in residual sum of squares (SSE) between the covariate-only model and the full model. Partial *R^2^* was then signed by the mean first derivative, so that an overall decrease gets a negative sign. Rate of change was derived by calculating the first derivative using the *gratia* package. This was done for all modalities.

To characterize temporal variations between the FA age-trajectories, we first divided nuclei age trajectories into those showing FA increase and those showing FA decrease. Only one nucleus (AV) fell into the decreasing group. To compare FA-increasing age trajectories according to their temporal profiles, we applied PCA on the normalized first derivative along the age trajectories, which reflects the rate of change at each time point. The resulting PC1 captures differences in the shape of the curvature of the trajectories, which contains information about the temporal properties (e.g. continuous increase versus early plateau). To characterise temporal patterns of whole-thalamus connections, the same approach was applied on all increasing FA trajectories (3 negative ones are excluded). PCA was also applied to the normalized first derivatives of functional connectivity and nucleus volume changes.

To compare nucleus-specific structural connection changes we generated a map comprising four clusters. Classification was based on the mean slope across the full age range, as well as the mean slope within the first and second halves of the age range. Connections were categorized as sustained high if they showed greater overall change than the average mean slope across nuclei, early dominant if they exhibited greater-than-average change during the first half of the age range, late dominant if they exhibited greater-than-average change during the second half of the age range, and sustained low if they showed relatively little change across the full age range.

### Thalamic feature maps for contextualization

#### Neuronal cell distribution

A key framework of the cell composition within the thalamus is the classification of core- and matrix cells. We used the map from (Müller et al., 2020), who used the Allen Human Brain Microarray data (Hawrylycz et al., 2012) to determine the relative expression of Parvalbumin (PV) and Calbindin (CB1), both calcium-binding proteins that are characteristic for thalamic core (PV) and matrix (CB1) cells. The ratio of PV and CB1 expression per voxel therefore provides a proxy for the core versus matrix cell proportion. The map is provided in MNI 152 space. To derive a core-matrix cell measure for each thalamic nucleus, we therefore used the above described consensus thalamus mask in MNI 152 space and extracted the average core-matrix standardized difference within each nucleus. Left and right hemisphere maps are derived from a different amount of donors (left 6, right 2) and the maps overlap to a different extent with the consensus mask. Therefore, to make our estimate per nucleus robust, we averaged the right and left hemisphere.

#### First order and higher order nucleus classification

A common classification of thalamic nuclei is the assignment of nuclei into first order nuclei, defined as getting ‘driver’ input from the subcortex and higher-order nuclei, defined as mostly receiving ‘driver’ input from cortex layer V. Here, we mainly adopt the classification proposed in (Sherman, 2005). An exception is made for nucleus AV, which was historically classified as a first order nucleus, however in a recent discussion it was stated as unclear (Perry and Mitchell, 2019). Therefore, we labeled AV as undefined.

#### SA-target area

The cortical SA axis has been defined in Sydnor *et al*. (2021) (Sydnor et al., 2021), by averaging rank orderings of ten different cortical feature maps encompassing micro- and macrostructural, functional, metabolic, transcriptomic, and evolutionary features (details described elsewhere (Sydnor et al., 2021), and describes an overarching cortical organization. Each Glasser parcel is assigned to a rank between 1 and 180, ordering the parcels along an axis spanning from sensorimotor to association regions. For each nucleus, we calculated the mean SA rank of the above defined nucleus-specific target parcels in structural and functional connectivity. This yielded in a SA score thalamic map for structural and for functional connectivity. Left and right hemisphere rank was averaged and yielded into a thalamic map of projected SA ranks.

### Associations between FA, thalamic feature maps, functional connectivity, and nucleus volume

#### Whole-thalamus patterns

The GAM derived features (i.e. partial *R*^2^, mean first derivative and PC1 score) of whole-thalamus structural connections were projected onto the cerebral cortex. Associations between those maps as well as the SA axis (Sydnor et al., 2021) were assessed using Spearman’s rank correlation and corrected for spatial autocorrelation using spin tests (Váša et al., 2018). We also accessed the relationship between this study’s age effect (partial *R*^2^) and the results of (Sydnor et al., 2025) (partial *R*^2^) by using Spearman’s rank correlation and spin tests.

#### Nucleus-specific patterns

To investigate the association between the temporal profiles (PC1) of thalamic nuclei and thalamic feature maps (core-matrix and SA target map), Spearman’s rank correlations were computed. Group differences in partial *R^2^* and PC1 between FO and HO nuclei were tested using Welch’s t-test, with effect size quantified using Hedges’ g. Akin, the relationship between magnitude of age effect (partial *R^2^*) and thalamic feature maps was assessed.

To examine the relation between structural and functional thalamocortical connectivity, SA scores derived from FA and functional connectivity were correlated (Spearman’s rank). For a first comparison of FA and functional connectivity partial *R^2^*, the normalized absolute difference between FA and functional connectivity partial *R^2^* was calculated for each nucleus. Finally, we calculated Spearman’s correlation of structural and functional connectivity partial *R^2^*for the full set of nuclei and a subset excluding LGN and MGN due to the early plateau in FA increase compared to the other nuclei. Further, the temporal profile components of FA and FC were Spearman correlated.

Last, to test the relationship of the development of nucleus volume development and nucleus-specific thalamocortical connections, Spearman rank correlations were again calculated for the full set and the subset of nuclei (LGN, MGN excluded) of FA partial *R^2^* and volume partial *R^2^*. The same was done for FC partial *R^2^* and volume partial *R^2^*. Further, the associations of the temporal profile components were tested using Spearman correlation.

## Data availability

The present work used existing data from the Human-Connectome Project Development (HCP-D, release 2.0) which is available via the NIMH Data Archive (https://nda.nih.gov/general-query.html?q=query=featured-datasets:HCP%20Aging%20and%20Development). For the contextualisation of our results, we used the following publicly available data: age effects of whole-thalamus structural connectivity changes in Glasser parcellation reported by Sydnor et al. (2025); the SA-axis (Sydnor et al., 2021) in Glasser parcellation, downloaded from https://github.com/PennLINC/S-A_ArchetypalAxis/tree/main/Glasser360_MMP); the core-matrix standardized difference map in MNI space (Müller et al., 2020), downloaded from https://github.com/macshine/corematrix; the classification of first- and higher order nuclei reported in Sherman (2005). The Glasser parcellation annotation file was downloaded from https://github.com/MICA-MNI/micapipe/tree/master/parcellations.

## Code availability

Custom code for all analysis pipelines will be made openly available on GitHub: https://github.com/John-Alexandra/TC_development. Preprocessing and tractography were performed using FSL, MRTRIX and Freesurfer, and data analyses were performed in Python and R. The segmentation of the thalamus was performed using the HIPS-THOMAS pipeline, available in https://github.com/thalamicseg/hipsthomasdocker. The modeling of age trajectories was performed in R (v. 4.5.0) and GAMS were fitted with the packages mgcv (v. 1.9-1, https://cran.r-project.org/src/contrib/Archive/mgcv/) and gratia (v. 0.10.0, https://cran.r-project.org/src/contrib/Archive/gratia/). Spin-test for parcellated cortical maps was performed using neuromaps.nulls.vasa (Váša et al., 2018, https://netneurolab.github.io/neuromaps/generated/neuromaps.nulls.vasa.html). Structural connections were visualized using the software braingl (https://github.com/rschurade/braingl). Thalamic nuclei were rendered using the python package pyvista (v. 0.46.1), brains surfaces were plotted using brainspace (v. 0.1.16), and network plots generated with pycirclize (v. 1.9.1). Additionally, we made use of the following python packages: seaborn (v. 0.13.2), colormaps (v. 0.4.2) (https://pratiman-91.github.io/colormaps/), matplotlib (v. 3.9.2), numpy (v. 2.0.2), pandas (v. 2.2.3), scipy (v. 1.13), nibabel (v. 5.3.2), neuromaps (v. 0.0.5), and scikit-learn (v. 1.5.2).

## Acknowledgements

A.J. was funded by Max Planck Society. A.S. was funded by the Max Planck Society. A.M. was funded by the German Academic Scholarship Foundation (Studienstiftung des deutschen Volkes) and the Max Planck Society’s Lise Meitner Excellence Program (awarded to SLV). J.R. was funded by Banting Fellowship. D.Y.E. was funded by Max Planck Society. V.J.S. was funded by T32MH016804 and T32MH018951 from the National Institute of Mental Health. B.W. is supported by funding from the Swiss National Science Foundation (grant number 32003B_219240). S.B.E. was funded by the European Union’s Horizon 2020 research and innovation programme (grant agreements 945539 [HBP SGA3], 826421 [VBC], and 101058516), the DFG (SFB 1451 and IRTG 2150), and the National Institute of Health (NIH; R01 MH074457). B.C.B. acknowledges research support from the National Science and Engineering Research Council of Canada (NSERC RGPIN-- 2025-- 05932), CIHR (FDN-- 154298, PJT-- 174995, PJT-- 191853, PJT-- 203761), SickKids Foundation (NI17-- 039), HIBALL, Healthy Brains and Healthy Lives (HBHL), Brain Canada Foundation, FRQS, the Tier-- 2 Canada Research Chairs Program, and the Centre for Excellence in Epilepsy at the Neuro (CEEN). A.W. was funded by Max Planck Society. S.L.V. was funded by the Max Planck Society through the Lise Meitner Excellence Program, the Jacobs Foundation Research Fellowship, the Hector Research Career Development Award, the European Research Council (ERC) Starting Grant, SOCO.

