## Supplementary Material for "Nucleus-level thalamic organization anchors multimodal signatures of thalamocortical maturation"

### This document includes:

**Supplementary Fig 1:** Age and sex distribution of HCP-D subsample.
**Supplementary Fig 2:** Group-level FA of whole-thalamus thalamocortical connections of left and right hemisphere.
**Supplementary Fig. 3:** Nucleus-specific target areas of structural connectivity left and right hemisphere.
**Supplementary Fig. 4:** FA age effects quantified by partial $R^2$.
**Supplementary Fig. 5:** Relation between FA age effect quantified by partial $R^2$ of structural nucleus-specific thalamocortical connections and thalamic and cortical features.
**Supplementary Fig. 6:** Nucleus-specific thalamocortical functional connectivity profiles of left and right hemisphere.
**Supplementary Fig. 7:** Age trajectories nucleus-specific thalamocortical functional connectivity.
**Supplementary Fig. 8:** Relation between PC1 of structural and functional age trajectories.
**Supplementary Fig. 9:** Age trajectories of whole-thalamus and nucleus volumes.
**Supplementary Fig. 10:** Temporal profile component of volume and association to temporal profile components of FA and FC.
**Supplementary Fig. 11:** Consistency thresholding and pruning of structural connections.

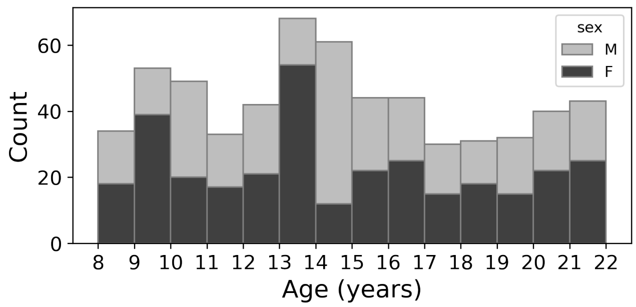


**Supplementary Fig. 1:** Age and sex distribution of HCP-D subsample used in this study (*N* = 604).

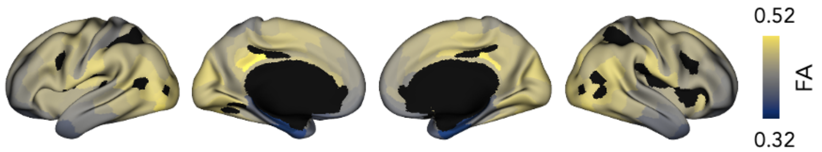


**Supplementary Fig. 2:** Group-level FA of whole-thalamus thalamocortical connections of left and right hemisphere. Black parcels indicate connections that were filtered out and midline.

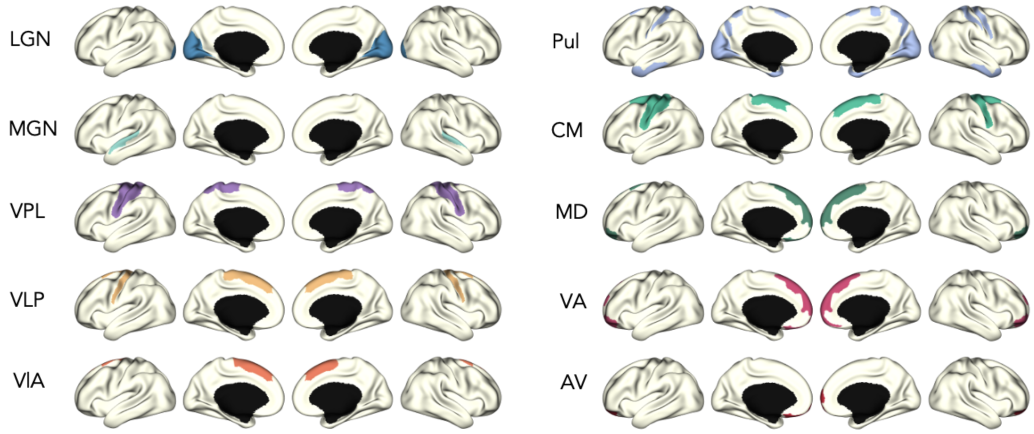


**Supplementary Fig. 3:** Nucleus-specific target areas of structural connectivity in left and right hemisphere.

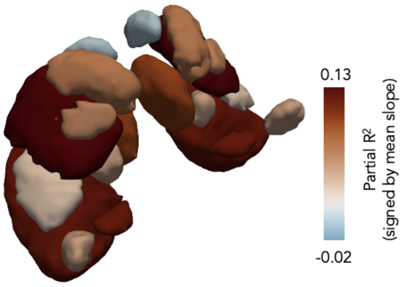


**Supplementary Fig. 4:** FA age effects quantified by partial $R^2$ (signed by mean slope) of nucleus-specific structural connections plotted on the thalamus.

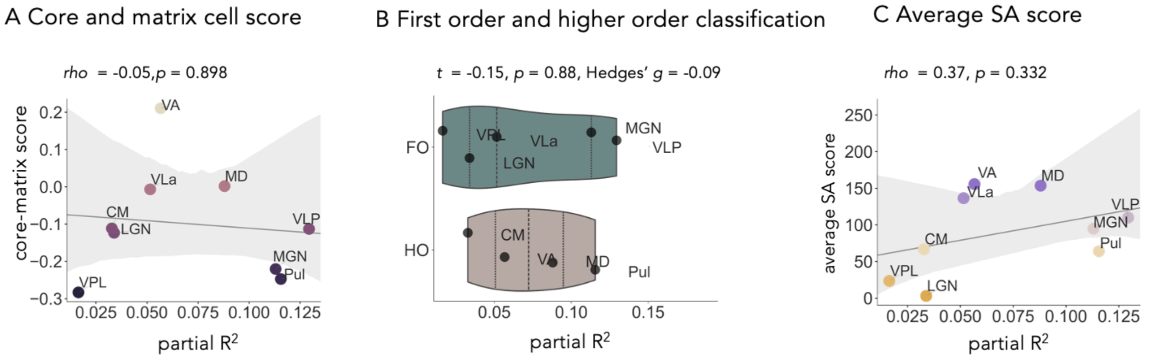


**Supplementary Fig. 5:** Relation between FA age effect quantified by partial $R^2$ (signed by mean slope) of structural nucleus-specific thalamocortical connections and thalamic and cortical features. **A** Scatterplot of partial $R^2$ and core-matrix score with a linear regression line and a 95% confidence interval. Scatter is colored by core-matrix score. **B** Violin plot of partial $R^2$ distribution in FO and HO. Results of t-test and Hedge's g are indicated. **C** Scatter plot of partial $R^2$ and SA scores with a linear regression line and a 95% confidence interval. Scatter is colored by SA scores. Results of Spearman's correlation and $p$ value are indicated.

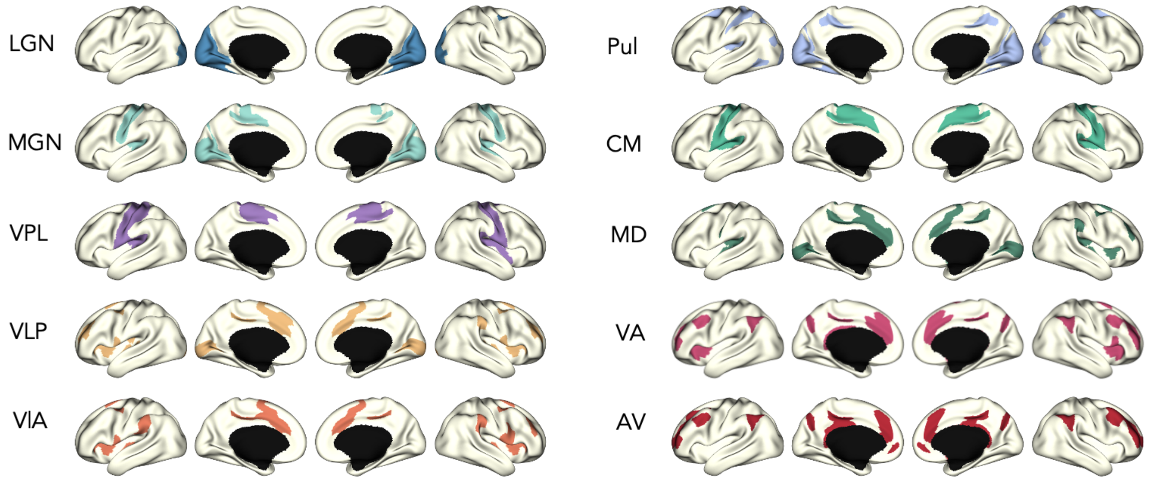


**Supplementary Fig. 6:** Nucleus-specific thalamocortical functional connectivity profiles of left and right hemisphere.

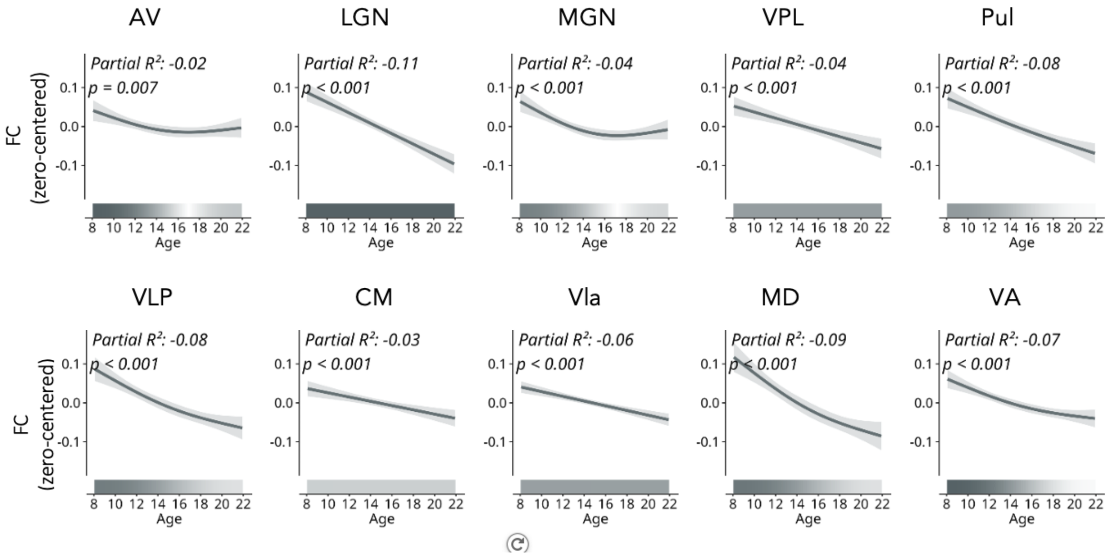


**Supplementary Fig. 7:** Age trajectories nucleus-specific thalamocortical functional connectivity. The trajectories represent zero-centered GAM smooth estimates with a shaded 95% confidence interval. Partial $R^2$ (signed by mean slope), and $p$-value (FDR-corrected) are indicated.

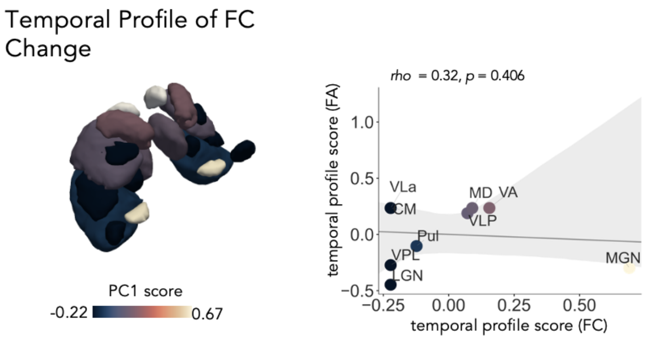


**Supplementary Fig. 8:** Relation between temporal profile scores of structural and functional age trajectories. PC1 scores of functional connectivity mapped onto thalamic nuclei. Scatter plot of PC1 of structural nucleus-specific thalamocortical connections and PC1 of functional nucleus-specific thalamocortical connections with a linear regression line and a 95% confidence interval.

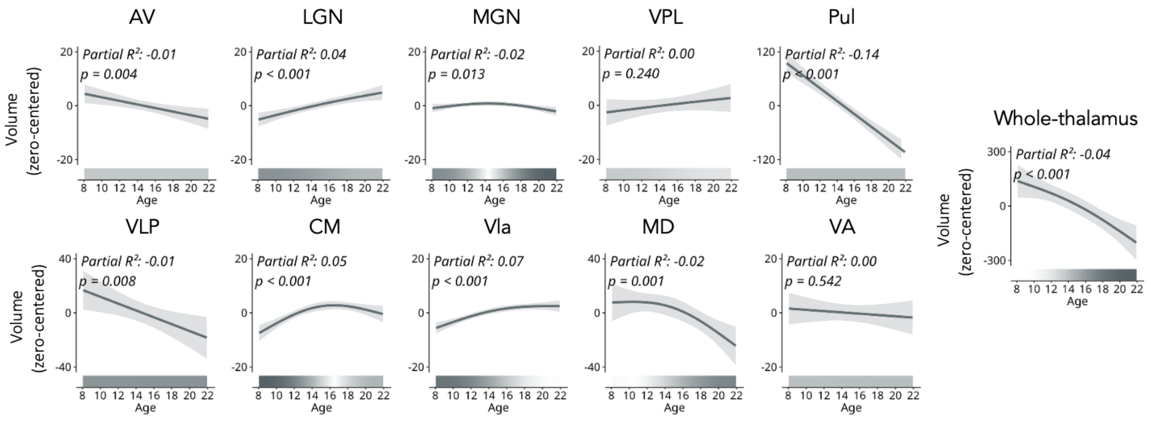


**Supplementary Fig. 9: Age trajectories of nucleus volumes and whole-thalamus.** The trajectories represent zero-centered GAM smooth estimates with a shaded 95% confidence interval. Partial $R^2$ (signed by mean slope), and $p$-value (FDR-corrected) are indicated.

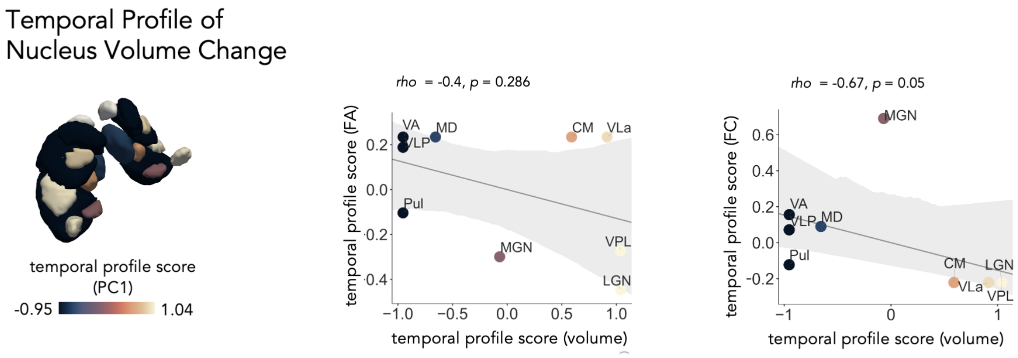


**Supplementary Fig. 10:** Temporal profile component of nucleus volume and association to temporal profile components of FA and FC. Principal component analysis (PCA) was applied to the normalized first derivatives sampled along the nucleus-volume trajectories and principal component 1 (PC1), referred to as temporal profile component, was plotted on the thalamus. Relation between temporal profile components of FA (left) and FC (right) and volume maturation with a linear regression line and a 95% confidence interval. Scatter is colored by volume-temporal profile components.

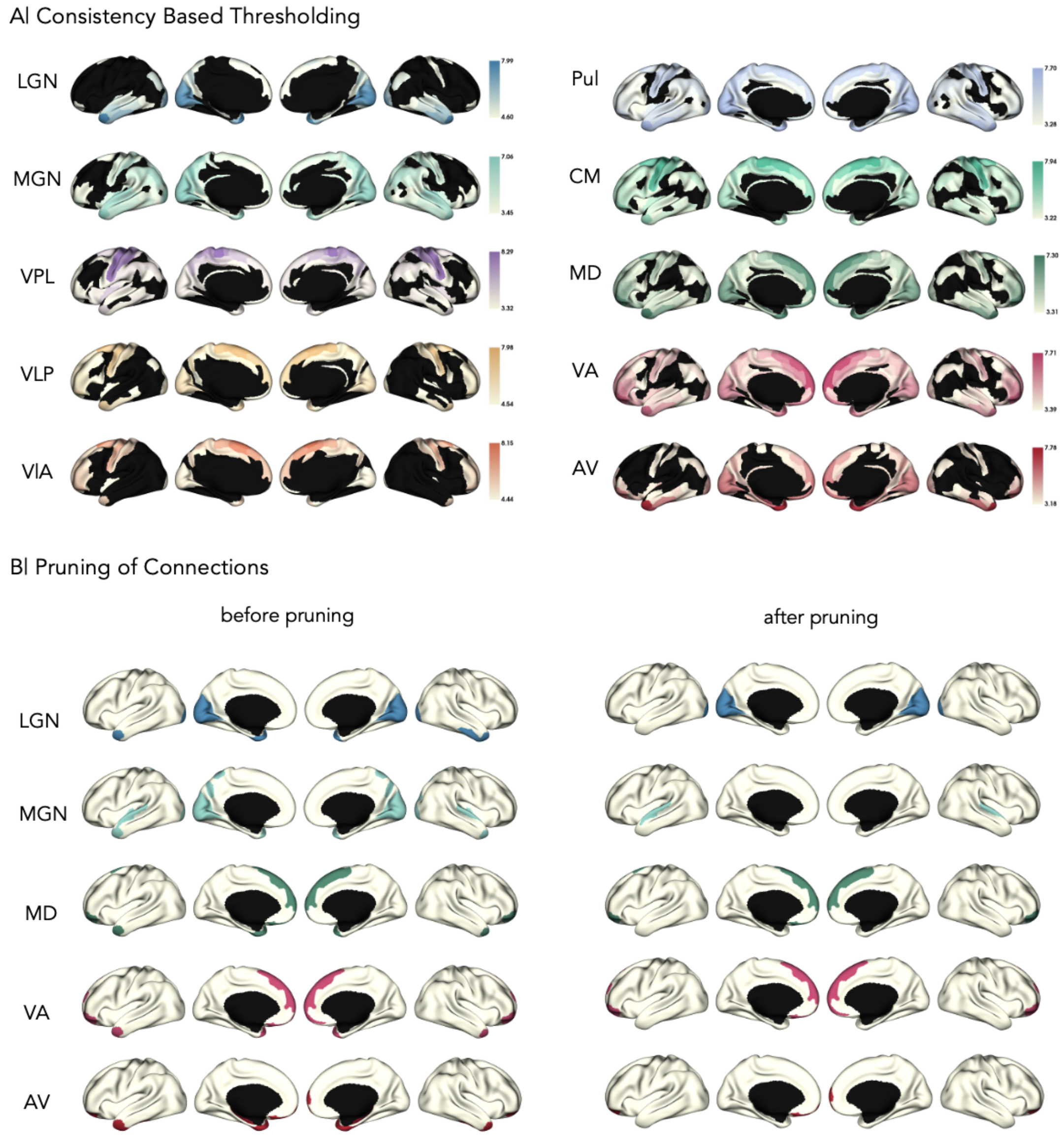


**Supplementary Fig. 11:** Consistency thresholding and pruning of structural connections. **A** Only connections present in at least 90% of participants retained (log-transformed relative streamline counts), whereas connections that did not match the criterion are masked out. **B** In 5 nuclei, on top of selection of strongest connections based on the top 10% using percentile-based thresholding (right), additional anatomical implausible connections were manually pruned out (left).
